# The engulfment receptor Draper is required for epidermal dendrite ensheathment

**DOI:** 10.1101/2025.09.18.677187

**Authors:** Chang Yin, Federico M. Tenedini, Amy Platenkamp, Jay Z. Parrish

**Affiliations:** Department of Biology, University of Washington, Campus Box 351800, Seattle, WA 98195, USA

## Abstract

Our experience of the external world is shaped by somatosensory neurons (SSNs) that innervate our skin and mediate responses to a range of environmental stimuli. The precise innervation patterns and response properties of SSNs are determined in part by specialized interactions with resident skin cells. One such interaction involves the preferential ensheathment of some SSNs by epidermal cells, an evolutionarily conserved intercellular interaction that regulates SSN morphogenesis and mechanical nociceptive sensitivity in *Drosophila*. The morphogenetic events during ensheathment resemble phagocytic engulfment, therefore we hypothesized that phagocytic receptors mediate molecular recognition of neurites to induce ensheathment. From a screen of epidermally expressed phagocytic receptors we found that the nimrod receptor gene *Draper* (*Drpr*) functions in epidermal cells to promote ensheathment. Endogenous Drpr accumulates at sites of epidermal ensheathment but not at epidermal contact sites with unensheathed neurites. Furthermore, overexpressing *Drpr* increased ensheathment selectively on neurons that are normally ensheathed, suggesting that molecular recognition by Drpr accounts for the specificity of ensheathment. Indeed, we found that an extracellular reporter for the Drpr ligand Phosphatidylserine (PS) accumulates at sites of ensheathment, and that preventing extracellular PS exposure by overexpressing the PS Flippase ATP8a blocked ensheathment. We additionally found that *Orion*, which encodes a chemokine-like protein that bridges Drpr-PS interactions, is required for sheath formation. Finally, we found that increasing ensheathment by overexpressing Drpr enhanced nociceptor sensitivity to mechanical stimulus. Altogether, these studies show that Drpr acts in epidermal cells to mediate molecular recognition events that drive ensheathment of neurites marked by extracellular PS.

## Introduction

An animal’s skin is densely innervated by somatosensory neurons (SSNs), which allow for perception and discrimination of touch, pressure, pain, and movement. Within the skin, many SSNs form specialized terminal structures that contribute to sensory transduction (Owens and Lumpkin, 2014; Zimmerman *et al*., 2014; Handler and Ginty, 2021; Yin *et al*., 2021). For example, afferent interactions with radially packed Schwann cell-derived lamellar cells in Pacinian corpuscles facilitate high frequency sensitivity (Nikolaev *et al*., 2020), and synapse-like contacts between mechanosensory Merkel cells and low threshold mechanoreceptors shape tactile inputs (Maricich *et al*., 2009; Woo *et al*., 2014; Hoffman *et al*., 2018). Likewise, anatomical studies dating back more than fifty years demonstrated that mammalian keratinocytes wrap free nerve endings including nociceptive C-fibers (Munger, 1965; Cauna, 1973, 1980), yet little is known about the molecular bases or functional relevance of keratinocyte-SSN coupling. However, epidermal cells in a range of invertebrates and vertebrates including *C. elegans*, *Drosophila* and *D. rerio* ensheath peripheral arbors of SSNs (Chalfie and Sulston, 1981; Han *et al*., 2012; Kim *et al*., 2012; O’Brien *et al*., 2012), providing tractable experimental systems to study this intercellular interaction. Of note, studies in *Drosophila* demonstrate that epidermal ensheathment influences SSN morphogenesis and sensitivity to noxious mechanical inputs (Jiang *et al*., 2014, 2019; Tenenbaum *et al*., 2017; Yang *et al*., 2019).

Epidermal sheath formation proceeds via a morphogenetic mechanism that is conserved from *Drosophila* to humans (Jiang *et al*., 2019; Talagas *et al*., 2020), with wrapping epidermal cells forming a mesaxon-like structure that is structurally similar to Schwann cell ensheathment of peripheral axons (Han *et al*., 2012; Kim *et al*., 2012; O’Brien *et al*., 2012). The signals that induce epidermal SSN ensheathment are unknown, but several features of the signaling system driving ensheathment have been characterized. First, neurites induce formation of epidermal sheaths: ensheathment is not observed in territory lacking neurites, and physical or genetic ablation of SSNs prevents sheath formation (O’Brien *et al*., 2012; Jiang *et al*., 2014, 2019). Only a subset of neurites that contact epidermal cells trigger ensheathment, suggesting that specific neurite signals are required to initiate the program rather than ensheathment being a default response to neurite contact. Second, neuronal signals are likely required for the maintenance of sheaths since ablation of ensheathed neurites triggers disassembly of epidermal sheaths (Jiang *et al*., 2019). Third, epidermal cells differentially ensheath subsets of SSNs, suggesting that different signals or different levels of a shared signal tune the levels of ensheathment in different neurons. In *Drosophila*, epidermal cells ensheath dendrite arborization (da) neurons, with nociceptive class IV da (C4da) neurons the most extensively ensheathed whereas proprioceptive C1da neurons exhibit minimal ensheathment (Jiang *et al*., 2019). Integrin-based ECM contacts in limit ensheathment of da neurons (Kim *et al*., 2012; Jiang *et al*., 2014), whereas heterophilic interactions between neuronal and epidermal isoforms of L1CAM/Neuroglian mediate dendrite-epidermis tethering to promote SSN ensheathment (Yang et al. 2019). However, the basis for selective recognition of target neurites for ensheathment by epidermal cells is not known.

Epidermal sheath formation shares hallmarks of the early stages of phagocytosis including local accumulation of Phosphatidylinositol 4,5-bisphosphate (PIP2), endocytic machinery, and an F-actin network at sites of nascent sheath formation (Scott *et al*., 2005). Additionally, epidermal cells act as primary phagocytes of neuronal debris with epidermally-expressed phagocytic receptors mediating the recognition events that drive epidermal neurite engulfment after injury and during pruning (Han *et al*., 2014; Rasmussen *et al*., 2015). We therefore hypothesized that receptors involved in phagocytic recognition may likewise mediate recognition events required for neurite ensheathment. The *Drosophila* genome contains ∼30 identifiable phagocytic receptors: 18 genes encoding scavenger receptors, which recognize an array of polyanionic ligands such as bacteria, apoptotic cell debris, and modified low-density lipoproteins; and 12 genes encoding Nimrod-type receptors, receptors for bacterial or apoptotic cell phagocytosis (Melcarne *et al*., 2019). We found that epidermal cells express a subset of these receptors, and that the Nimrod family receptor Draper (Drpr) is required in epidermal cells for sheath formation. Furthermore, epidermal *Drpr* overexpression increased ensheathment selectively on neurons that are normally ensheathed and enhanced the sensitivity of nociceptors to mechanical stimuli. Consistent with the model that Drpr mediates recognition events that drive ensheathment, extracellular reporters for the Drpr ligand phosphatidylserine (PS) accumulate at sites of ensheathment, blocking dendritic PS exposure prevented ensheathment, and the PS bridging molecule Orion was required for sheath formation. Altogether these results support a model in which epidermally expressed Drpr recognizes extracellular PS on dendrites to initiate epidermal sheath formation.

## Results

### Engulfment receptors regulate multiple aspects of epidermal dendrite ensheathment

We used RNA sequencing (RNA-Seq) of acutely dissociated larval epidermal cells (Yoshino *et al*., 2025) to define the repertoire of epidermally-expressed engulfment receptors. To validate our ability to detect biologically relevant levels of engulfment receptor expression we additionally monitored engulfment receptor expression in larval hemocytes, *Drosophila* immune cells with a broad range of phagocytic activities (Melcarne *et al*., 2019). We isolated larval body wall skin preparations, dissociated them into single cell suspensions and manually isolated GFP-labeled epidermal cells (*R38F11-GAL4, UAS-2xEGFP*) or hemocytes (*Hml-GAL4, UAS-2xEGFP*) for SMRT-seq gene expression analysis (Picelli *et al*., 2014). In hemocytes we reproducibly detected expression of the majority (27/30) of *Drosophila* engulfment receptor genes, 17 of which were expressed at levels > 1 transcript per million (TPM; Figure 1A). In contrast, epidermal cells expressed a more limited repertoire of receptors that was on average expressed at lower levels (2/16 expressed at levels > 1 TPM). The most highly expressed epidermal phagocytic receptor genes include *Crq* and *Drpr* which participate in epidermal phagocytosis of damaged dendrites (Han *et al*., 2014), but epidermal cells additionally expressed several receptors with no previously characterized function in epidermal phagocytosis.

**Figure 1.**
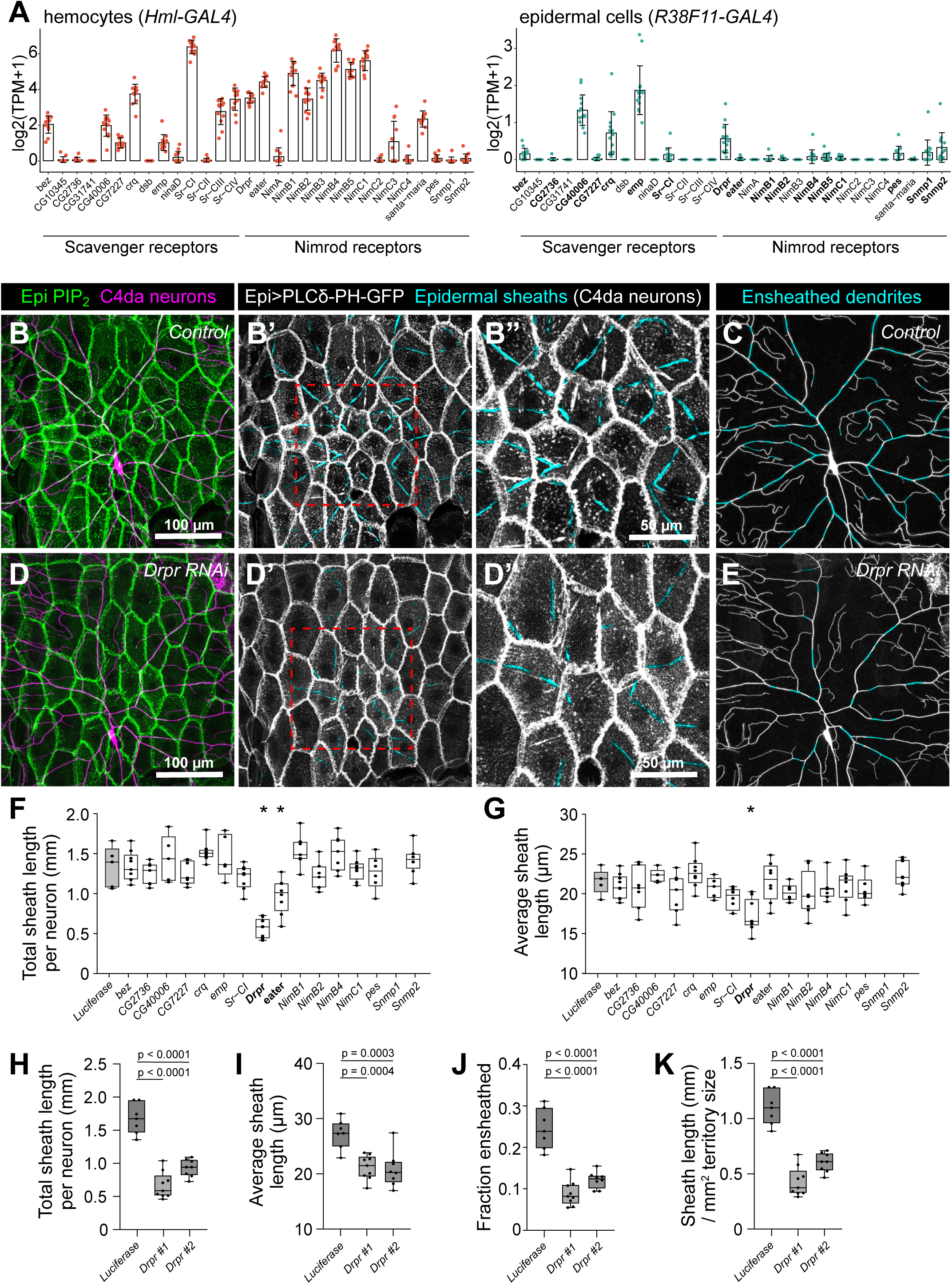
The nimrod receptor Drpr regulates epidermal dendrite ensheathment. (A) RNA-seq analysis of scavenger and nimrod family receptor gene expression in hemocytes and epidermal cells. Each point represents an independent biological sample derived from a library of 10 purified GFP-expressing cells. TPM, transcripts per million. Epidermal candidate genes in bold exhibited a TPM value > 0.1 in at least one biological sample and were selected for further analysis. Genotypes: *Hml-GAL4, UAS-2xEGFP / +* (hemocytes); *R38F11-GAL4, UAS-2xEGFP / +* (epidermal cells). (B-G) Genetic screen for phagocytic receptors that control epidermal dendrite ensheathment. (B-E) Dual labeling of epidermal sheaths (*A58-GAL4, UAS-PLC^δ^-PH-GFP*) and nociceptive C4da neurons (*ppk-CD4-tdTomato*) in third instar larvae expressing control (*Luciferase*) or *Drpr* RNAi transgenes in epidermal cells. Maximum intensity projections of confocal stacks show the distribution of C4da dendrites over epidermal cells in a single dorsal hemisegment. Composite images depict the distribution of epidermal sheaths labeled by the PIP2 reporter PLC^δ^-PH-GFP along C4da arbors. Genotypes: *A58-GAL4, UAS-PLC^δ^-PH-GFP / UAS-Luciferase-RNAi* (control) or *A58-GAL4, UAS-PLC^δ^-PH-GFP / UAS-Drpr-RNAi*. (F-G) Quantification of screen results. Plots depict the total sheath length per neuron (F) and the average length of individual sheaths (G). Box plots here and in subsequent panels depict mean values and 1st/3rd quartile, whiskers mark maximum and minimum values, and individual data points are shown (n;: 5 neurons per genotype). *P<0.05 compared to *Luciferase RNAi* control; one-way ANOVA with post-hoc Dunnett’s multiple comparisons test. (H-K) Valdiation of *Drpr* knockdown phenotypes. Plots depict the total length of epidermal sheaths per neuron (H), the average length of individual sheaths (I), the proportion of C4d dendrite arbors ensheathed in PLC-PH-GFP-positive epidermal membrane domains (J), and the total sheath length normalized to territory size for larvae expressing control (*Luciferase*) or each of two distinct *Drpr* RNAi transgenes in epidermal cells. n = 8 neurons per genotype. P values, one-way ANOVA with post-hoc Dunnett’s multiple comparisons test. <u>Experimental Genotypes:</u> *(A) *<u>hemocytes</u>*: w^1118^; Hml-GAL4, UAS-2xEGFP / +* *<u>epidermal cells</u>*: *w^1118^; R38F11-GAL4, UAS-2xEGFP / +* *(B-E) *<u>Control</u>*: w^1118^; ppk-CD4-tdTomato^4A^ / +; A58-GAL4, UAS-PLC^δ^-PH-GFP, ppk-CD4-tdTomato^10A^ / UAS-Luciferase-RNAi* *<u>Drpr RNAi</u>*: *w^1118^; ppk-CD4-tdTomato^4A^ / +; A58-GAL4, UAS-PLC^δ^-PH-GFP*, *ppk-CD4-tdTomato^10A^ / UAS-Drpr-RNAi^HMS01623^* *(F-G) *<u>RNAi screen:</u>* w^1118^; ppk-CD4-tdTomato^4A^ / +; A58-GAL4, UAS-PLC^δ^-PH-GFP,* *ppk-CD4-tdTomato^10A^ / UAS-RNAi transgene* *(H-K) <u>Control</u>: w^1118^; ppk-CD4-tdTomato^4A^ / +; A58-GAL4, UAS-PLC^δ^-PH-GFP, ppk-CD4-tdTomato^10A^ / UAS-Luciferase-RNAi* *<u>Drpr RNAi 1</u>*: *w^1118^; ppk-CD4-tdTomato^4A^ / +; A58-GAL4, UAS-PLC^δ^-PH-GFP*, *ppk-CD4-tdTomato^10A^ / UAS-Drpr-RNAi^HMS01623^* *<u>Drpr RNAi 2</u>*: *w^1118^; ppk-CD4-tdTomato^4A^ / +; A58-GAL4, UAS-PLC^δ^-PH-GFP, ppk-CD4-tdTomato^10A^ / UAS-Drpr-RNAi^HMS01623^*

To assay contributions of epidermal engulfment receptor genes to dendrite ensheathment we used RNAi to knock down receptor gene expression selectively in epidermal cells that additionally expressed the phosphatidylinositol 4,5-bisphosphate (PIP_2_) reporter PLCδ-PH-GFP (Várnai and Balla, 1998) to label epidermal sheaths. PIP2 accumulation on epidermal membranes is the earliest identified marker of epidermal sheaths, and PIP2 labeling persists in mature sheaths (Jiang *et al*., 2019), hence we monitored for alterations in the extent of PIP2 labeling of epidermal membrane stretches adjacent to C4da neurons, which are extensively ensheathed (Jiang *et al*., 2014, 2018, 2019). First, we assayed for ensheathment in larvae expressing a control RNAi transgene (*R38F11-GAL4, UAS-Luc-RNAi*) and found that ∼1.45 mm of the C4da dendrite arbor was ensheathed by PLCδ-PH-GFP-positive stretches of epidermal membrane that were ∼22 μm in length, on average (Figure 1, B to E). Similarly, epidermal RNAi of 14/16 engulfment receptor genes had no significant effect on the extent of dendrite ensheathment in third instar larvae (Figure 1, F and G), consistent with the notion that most epidermal engulfment receptors are dispensable for dendrite ensheathment. In contrast, RNAi of *Drpr* and to a lesser extent *ea*, two engulfment receptor genes required for epidermal phagocytosis of damaged dendrites (Han *et al*., 2014), caused significant reductions in dendrite ensheathment (Figure 1, D to G). Hence, phagocytic receptors required for engulfment of damaged dendrites are also required for epidermal dendrite ensheathment.

Epidermal sheath formation is restricted to sites of neurite contact and epidermal sheaths are disassembled following dendrite regression (Jiang *et al*., 2019), suggesting that short-range neuron-derived signals are required for both establishment and maintenance of epidermal sheaths. We therefore examined whether epidermal phagocytic receptors were involved in maintaining the coupling of neurites and epidermal sheaths, using “empty” sheaths as a proxy for exuberant sheath extension or defects in sheath disassembly following dendrite regression. In control larvae, we rarely observed empty sheaths: we identified an average of 65.6 ± 9.9 PLCδ-PH-GFP positive sheath structures per C4da neuron, but only 0.9 ± 0.7 of these lacked complete dendrite innervation (Supplemental Figure S1). Similarly, RNAi of *Drpr* or *ea* in epidermal cells induced no significant increase in empty sheaths. However, RNAi of a single receptor gene, the nimrod family receptor *NimB4*, led to a significant increase in sheath structures lacking neurites (Supplemental Figure S1). Hence, distinct epidermal phagocytic receptors regulate different aspects of neurite ensheathment that may involve different recognition events. For the remainder of this study, we focused on characterizing *Drpr* function in epidermal dendrite sheath formation.

### Drpr functions in epidermal cells to promote epidermal sheath formation

To further establish the requirement for *Drpr* in epidermal dendrite ensheathment, we assayed effects of epidermal *Drpr* RNAi using a second RNAi transgene that targets a different portion of the *Drpr* mRNA (MacDonald *et al*., 2006). As with the first transgene, expression of the second *Drpr* RNAi transgene induced significant reductions in the overall and average length of epidermal sheaths (Figure 1, H and I), demonstrating that the observed ensheathment phenotypes are caused by *Drpr* knockdown and not off-target effects. Next, to determine whether the observed ensheathment defects reflected a secondary consequence of dendrite branching defects or a primary effect on ensheathment we normalized the epidermal sheath length measurements to dendrite length and territory size, respectively. Indeed, we found that epidermal *Drpr* knockdown caused a significant reduction in ensheathment that is independent of effects on dendrite arborization or territory size (Figure 1, J and K).

Formation of PIP_2_-enriched microdomains on epidermal membranes adjacent to sensory neurites is followed by recruitment of the GTPase Rho1 and filamentous actin (F-actin) and subsequently junctional proteins, which may seal sheaths and limit sheath permeability (Kim *et al*., 2012; Jiang *et al*., 2019). *Drosophila* sheaths contain adherens junction and numerous septate junction proteins including Band4.1/coracle (cora) (Kim *et al*., 2012; Tenenbaum *et al*., 2017; Jiang *et al*., 2019; Yang *et al*., 2019), therefore to further corroborate the requirement for *Drpr* function in dendrite ensheathment we assayed for effects of an amorphic *Drpr* allele (*Drpr^/15^*) on cora accumulation at epidermal sheaths. Compared to wild-type controls, larvae homozygous for *Drpr^/15^* exhibited significant reductions in the number and total length of cora-positive membrane domains adjacent to C4da dendrites (Figure 2, A to D). Furthermore, the cora-positive sheaths that formed in *Drpr^/15^* mutants exhibited significant reductions in diameter (Figure 2E).

**Figure 2.**
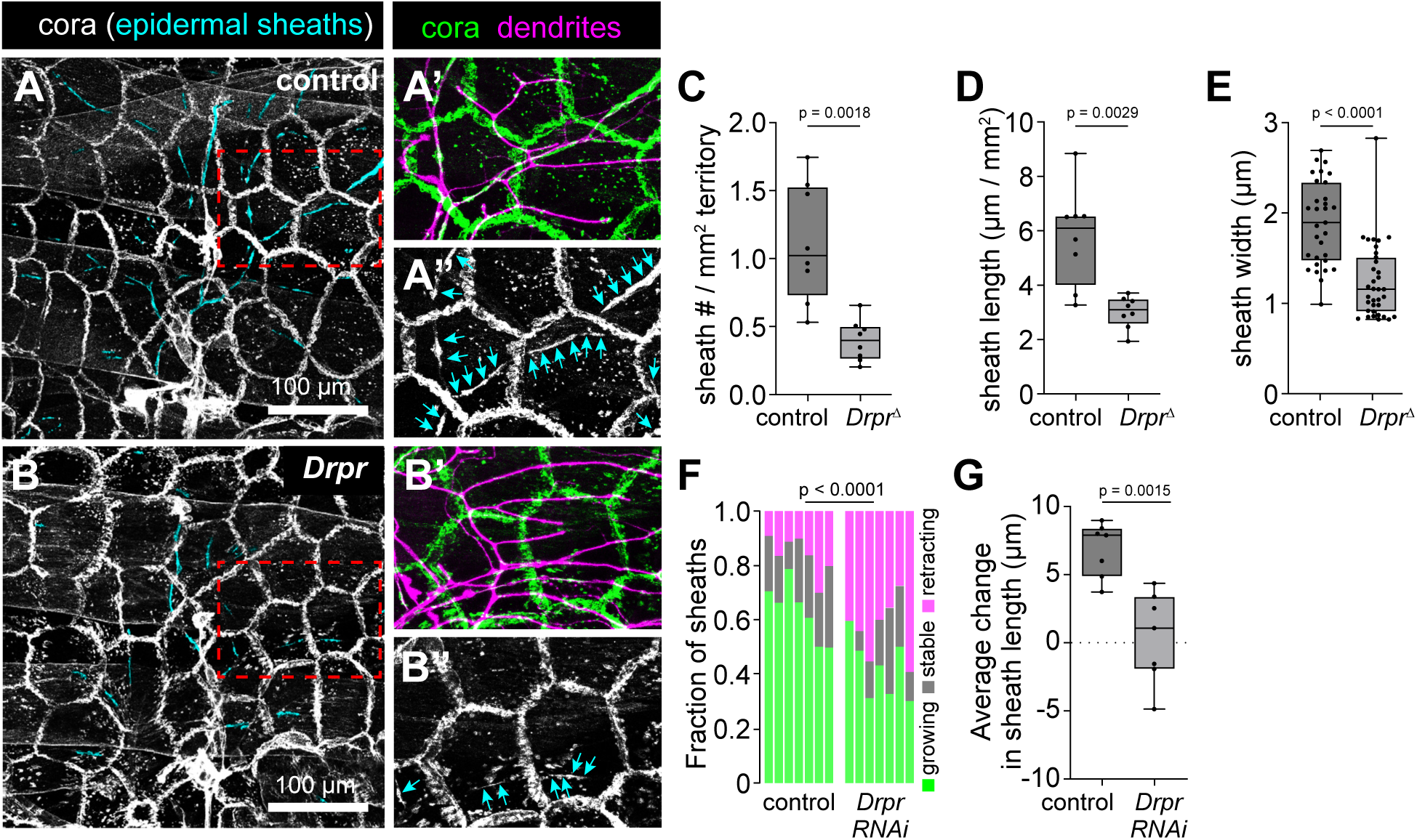
***Drpr* is required in epidermal cells for dendrite ensheathment**. (A-E) Drpr is required for formation of mature epidermal sheaths. (A-B) Maximum intensity projections show the distribution of cora immunoreactivity in the epidermis of (A) wild-type control and (B) *Drpr^/1^* amorphic mutant larvae. Cora accumulates at intercellular junctions and at epidermal sheaths, which are pseudocolored cyan. Zoomed images show double labeling of dendrites with anti-HRP antibodies (magenta) and cora (A’ and B’), with epidermal sheaths marked by cyan arrows (A” and B”). Plots depict the total number (C) and length (D) of cora+ sheaths per neuron normalized to the two-dimensional area covered by the dendrite arbor, and (E) the mean width of individual epidermal sheaths. n = 9 neurons each genotype for (C, D); 31 sheaths from control and 35 sheaths from *Drpr* mutants (from 6 neurons each) were analyzed in (E). *P<0.05 compared to controls, unpaired t-test with Welch’s correction (C and D) or Kolmogorov-Smirnov test (E). (F-G) *Drpr* knockdown alters sheath dynamics. (F) Plot depicts the distribution of dynamic events (growth, retraction, stable sheaths) over a 12 h time lapse for controls or larvae expressing epidermal *Drpr* RNAi. *Drpr* RNAi larvae exhibit significantly fewer growth events and more retraction events than controls. n = 7 neurons each genotype; 445 total sheaths for control, 281 for *Drpr* RNAi.. ***P < 0.001, Chi-square test. (G) Plot depicts the mean change in sheath length for controls or larvae expressing epidermal *Drpr* RNAi. n = 7 neurons each genotype. P value, unpaired t-test with Welch’s correction. (A-E) *<u>Control</u>*: w^1118^ *<u>Drpr</u>: w^1118^;; Drpr^/15^* (F-G) *<u>Control</u>: w^1118^; ppk-CD4-tdTomato^4A^ / +; A58-GAL4, UAS-PLC^δ^-PH-GFP, ppk-CD4-tdTomato^10A^ / UAS-Luciferase-RNAi* *<u>Drpr</u>: w^1118^; ppk-CD4-tdTomato^4A^ / +; A58-GAL4, UAS-PLC^δ^-PH-GFP, ppk-CD4-tdTomato^10A^ / UAS-Drpr-RNAi^HMS01623^*

To gain insight into the cellular origin of the ensheathment defects caused by *Drpr* inactivation we used time-lapse imaging to monitor effects of epidermal *Drpr* RNAi on sheath dynamics. Over a 12 h time-lapse during the time window when sheaths normally form (96-108 h AEL), we identified two principal differences in growth behavior of PLC^δ^-PH-GFP-positive epidermal sheaths in control and *Drpr* RNAi larvae. First, epidermal *Drpr* RNAi altered the frequency of dynamic events, increasing retraction events and reducing growth events (Figure 2F). Nevertheless, most sheaths in *Drpr* RNAi larvae were stable or growing during the time lapse (Figure 2F) suggests that *Drpr* is dispensable for maintenance of existing sheaths. Second, *Drpr* RNAi significantly reduced the extent of sheath growth; individual sheaths grew by∼8 uM in controls and only ∼1 uM in *Drpr* RNAi larvae (Figure 2G). Sheaths continued to form, albeit at reduced rates in *Drpr* RNAi larvae, consistent with the model that Drpr acts partially redundantly with an additional receptor(s) to mediate neurite recognition and drive ensheathment.

### Drpr is enriched at epidermal sheaths but not at other dendrite-epidermis contact sites

Dendrites of all da neurons grow in contact with epidermal cells, but distinct classes of da neurons have different capacities to induce epidermal sheath formation (Jiang *et al*., 2019). We reasoned that if Drpr is involved in recognition events that drive ensheathment then Drpr should coalesce at sites of ensheathment but not contact sites with unensheathed dendrites. As a first test of this model, we examined whether endogenous Drpr protein accumulated at sites labeled by the ensheathment marker PLCδ-PH-GFP. Indeed, Drpr immunoreactivity was enriched at PIP2-positive membrane domains in epidermal cells, both at epidermal intercellular junctions and at epidermal sheaths, with >80% of PIP-2-positive sheaths co-labeled by Drpr immunoreactivity (Drpr+) (Figure 3, A and B). We likewise found that Drpr immunoreactivity was enriched at mature epidermal sheaths, which are labeled by antibodies to cora: >70% of cora-positive epidermal sheaths also containing stretches of Drpr immunoreactivity (Figure 3, C and D). In contrast, anti-Drpr immunoreactivity at cora-positive membrane domains was eliminated in *Drpr* mutant larvae (Supplemental Figure 2), demonstrating specificity of the antibody and suggesting that the incomplete loss of ensheathment in *Drpr* mutants reflects contributions of a redundant pathway rather than protein perdurance. Hence, Drpr accumulates at sites of dendrite ensheathment and persists in mature sheaths.

**Figure 3.**
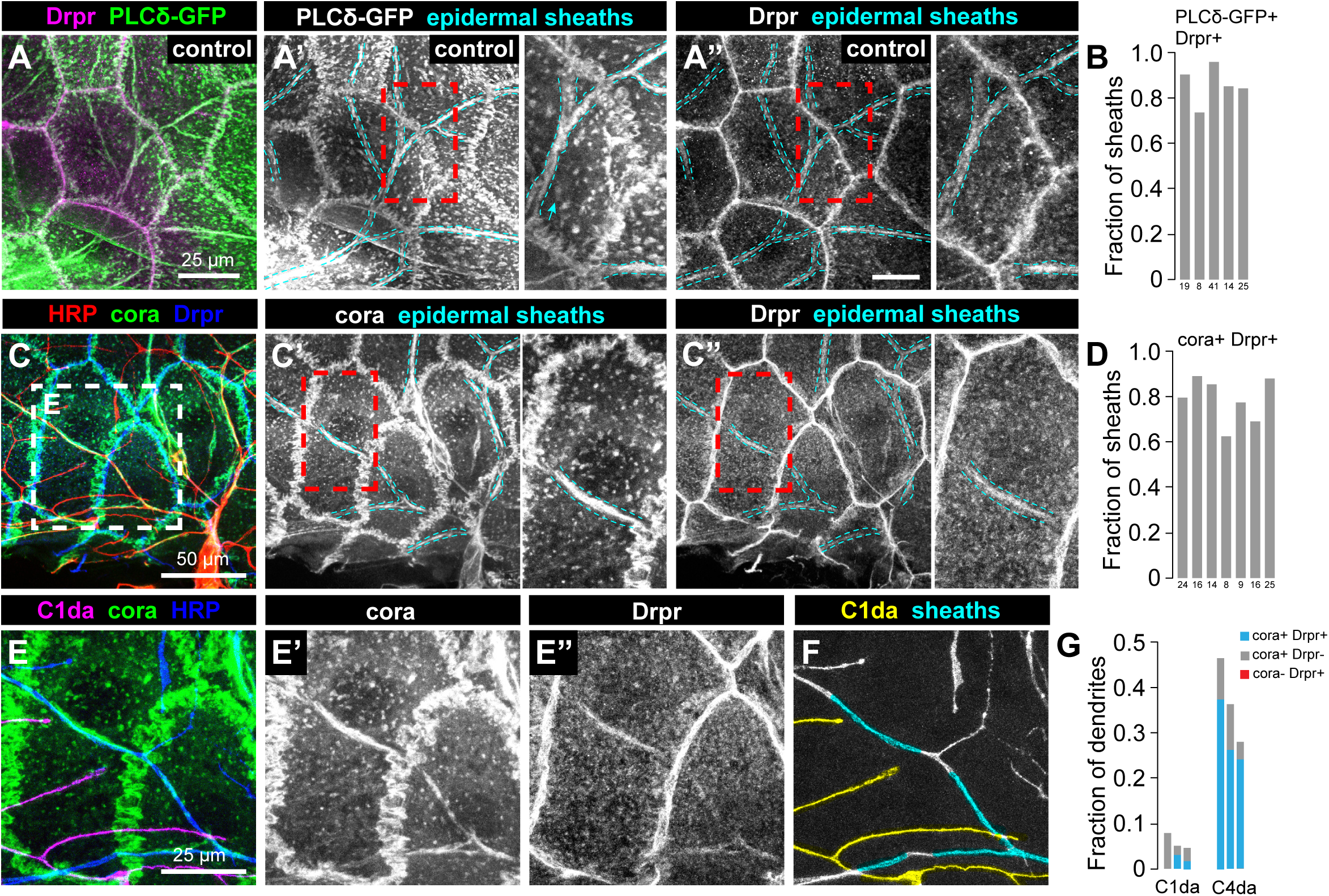
Drpr is enriched at epidermal sheaths but not other sites of dendrite-epidermis contact. (A-B) Drpr localizes to epidermal sheaths. (A) Composite image shows immunoreactivity a larval fillet preparations double-labeled for the PIP2 reporter PLC^δ^-PH-GFP and Drpr. Cyan hatched lines in region of interest (red box) outline PIP2-positive epidermal sheaths and sheath-localized Drpr immunoreactivity. (B) Plot depicts the proportion of PIP2-positive sheaths that were labeled by anti-Drpr immunoreactivity. (C-D) Drpr persists at mature epidermal sheaths. (C) Composite image shows immunoreacitivy for cora, a marker of mature sheaths, HRP to label sensory neurons and Drpr. Blue arrows mark cora+ epidermal sheaths. (D) Plot depicts the proportion of cora+ epidermal sheaths that were labeled by anti-Drpr immunoreactivity. (E-F) Drpr accumulates at sites of ensheathment but not at epidermal contact sites with unensheathed neurites. (E) Composite image shows immunoreacitivy for cora, HRP, and Drpr. Dendrites of C1da neurons are pseudocolored magenta in (E) and yellow in (F), and cora+ sheathes are pseudocolored cyan in (F). (G) Drpr accumulates at cora+ sheaths, but not at cora-stretches in C1da or C4da neurons. Bars depict the proportion of dendrite arbors from C1da and C4da neurons in contact with cora+ Drpr+ (blue), cora+ Drpr-(gray), and cora-Drpr+ (red) epidermal membrane domains. <u>Experimental Genotypes:</u> (A-B) *<u>Control</u>: w^1118^;; A58-GAL4, UAS-PLC^δ^-PH-GFP / +* (C-E) *<u>Control</u>: w^1118^* (F-G) *<u>Control</u>: w^1118^; 98b-GAL4, UAS-CD4-tdGFP^8M2^*

Next, we examined whether Drpr selectively accumulates at sites of dendrite ensheathment or more broadly accumulates at sites of dendrite contact by assaying for Drpr accumulation at epidermal contact sites with C1da dendrites, which are minimally ensheathed. To this end, we stained larval fillet preparations with antibodies Drpr in combination with antibodies to cora to label epidermal sheaths and HRP to label sensory dendrites. We found that Drpr+ stretches were present only at sites additionally labeled by the sheath marker cora, whereas Drpr+ stretches were absent from epidermal contact sites with dendrites of C1da neurons or from unensheathed portions of C4da dendrite arbors (Figure 3, E to G). Hence, Drpr accumulation at sites of dendrite contact is selective for sites of ensheathment.

### The engulfment receptor Drpr regulates dendrite branching

Blocking epidermal ensheathment alters dendrite morphogenesis of ensheathed but not unensheathed neurons (Parrish *et al*., 2009; Jiang *et al*., 2014, 2019), therefore we next investigated effects of *Drpr* inactivation on dendrite branching in C1da neurons and C4da neurons. Compared to wild-type controls, *Drpr^/15^* mutation had no significant effect on dendrite branch number or total dendrite length in C1da neurons (Figure 4, A to C). In contrast, we found that *Drpr^/15^* mutant larvae exhibited a significant increase in the overall density of body wall innervation by C4da dendrites (Figure 4, D and E). *Drpr* mutant larvae exhibited a significant increase in terminal dendrite branch number, particularly short terminal (< 20 μm) branches (Figure 4, F and G), similar to other treatments that block epidermal dendrite ensheathment (Parrish *et al*., 2009; Jiang *et al*., 2014, 2019; Tenenbaum *et al*., 2017). *Drpr* is therefore required to limit terminal branching in neurons with ensheathed but not unensheathed dendrites.

**Figure 4.**
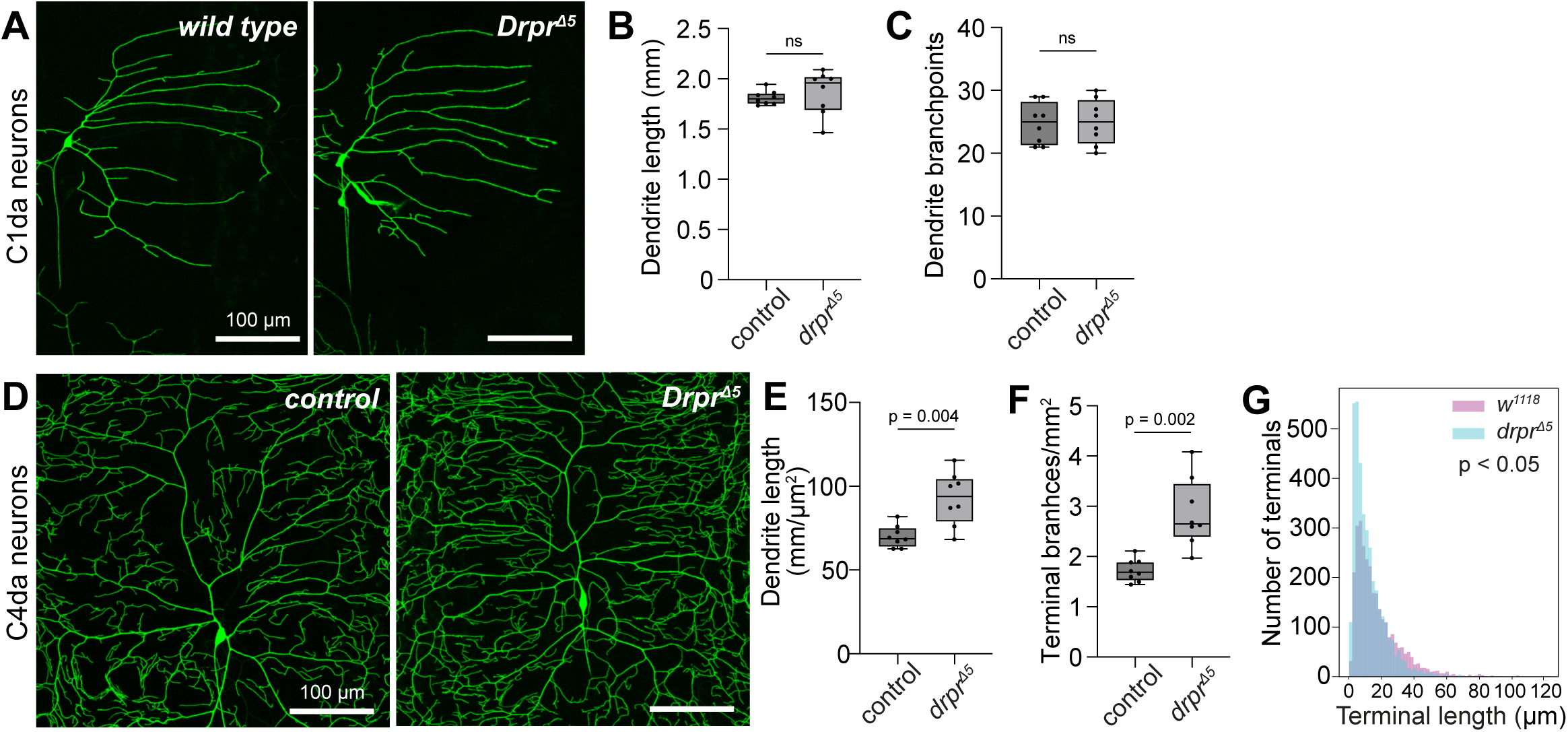
*Drpr* regulates morphogenesis of ensheathed but not unensheathed dendrite arbors. (A-C) *Drpr* is dispensable for denrite morphogenesis in C1da neurons, whose arbors exhibit minimal ensheathment. (A) Maximum intensity projections from wild-type control and *Drpr* mutant larvae depict C1da neurons expressing *CD4-tdGFP* (*98b-GAL4*). Plots depict (B) dendrite length normalized to receptive field size and (C) the number of dendrite branchpoints per C1da neuron. n = 8 neurons for each genotype. Comparing values from control and *Drpr* mutant larvae using a Mann-Whitney test revealed no significant differences. (D-G) *Drpr* restricts terminal dendrite branching in C4da neurons, which are extensively ensheathed. (D) Maximum intensity projections show dendrite arbors of representative C4da neurons labeled by *ppk-CD4-tdGFP* in wild-type control and *Drpr* mutant larvae. C4da neurons from *Drpr* mutants exhibited a significant increase in dendrite density (E) and terminal dendrite number (F). n = 8 neurons for each genotype. *P<0.05 compared to wild-type controls, Mann-Whitney test. (G) Histogram depicts the distribution of terminal branch lengths from wild-type control and *Drpr* mutant C4da neurons. *P<0.05 compared to wild-type controls, Kolmogorov–Smirnov Two-Sample Test sampling 521 terminal dendrites from *Drpr* mutants and 1198 terminal dendrites from controls. Sampling equally sized random subsets from each yielded a similar result. <u>Experimental Genotypes:</u> (A-C) *<u>Control</u>: w^1118^; 98b-GAL4, UAS-CD4-tdGFP^8M2^* *Drpr^/15^: w^1118^; 98b-GAL4, UAS-CD4-tdGFP^8M2^; Drpr^/15^* (D-G) *<u>wild type</u>: w^1118^;; ppk-CD4-tdTomato^10A^ Drpr^/15^*: *w^1118^;; Drpr^/15^, ppk-CD4-tdTomato^10A^*

### Drpr overexpression induces exuberant ensheathment activity

Based on our findings that *Drpr* is required for formation of epidermal dendrite sheaths, Drpr is enriched at sites of ensheathment but not other sites of dendrite-epidermis contact, and Drpr co-localizes with the early ensheathment marker PLCδ-PH-GFP, we hypothesized that Drpr mediates dendrite recognition to trigger epidermal ensheathment. To further define Drpr function in dendrite recognition we examined whether epidermal *Drpr* overexpressed was sufficient to induce ectopic dendrite ensheathment. First, we assessed effects of *Drpr* overexpression on ensheathment of C4da dendrites, the most extensively ensheathed among da neurons. Using two independent markers of epidermal sheaths, we found that epidermal *Drpr* overexpression (*R38F11-GAL4, UAS-Drpr*) significantly increased the overall proportion of C4da dendrites that were ensheathed compared to effector-only controls (*UAS-Drpr*) (Figure 5, A-D, and Supplemental Figure 3). In contrast, epidermal *Drpr* overexpression did not induce ectopic sheath formation on C1da dendrites, which normally exhibit minimal ensheathment (Figure 5, E to G). Hence, *Drpr* overexpression is sufficient to trigger exuberant ensheathment, but the selectivity for preferred target(s) is maintained. These results further suggest that the extent of ensheathment is not strictly determined by availability of neuronal signals required for epidermal recognition.

**Figure 5.**
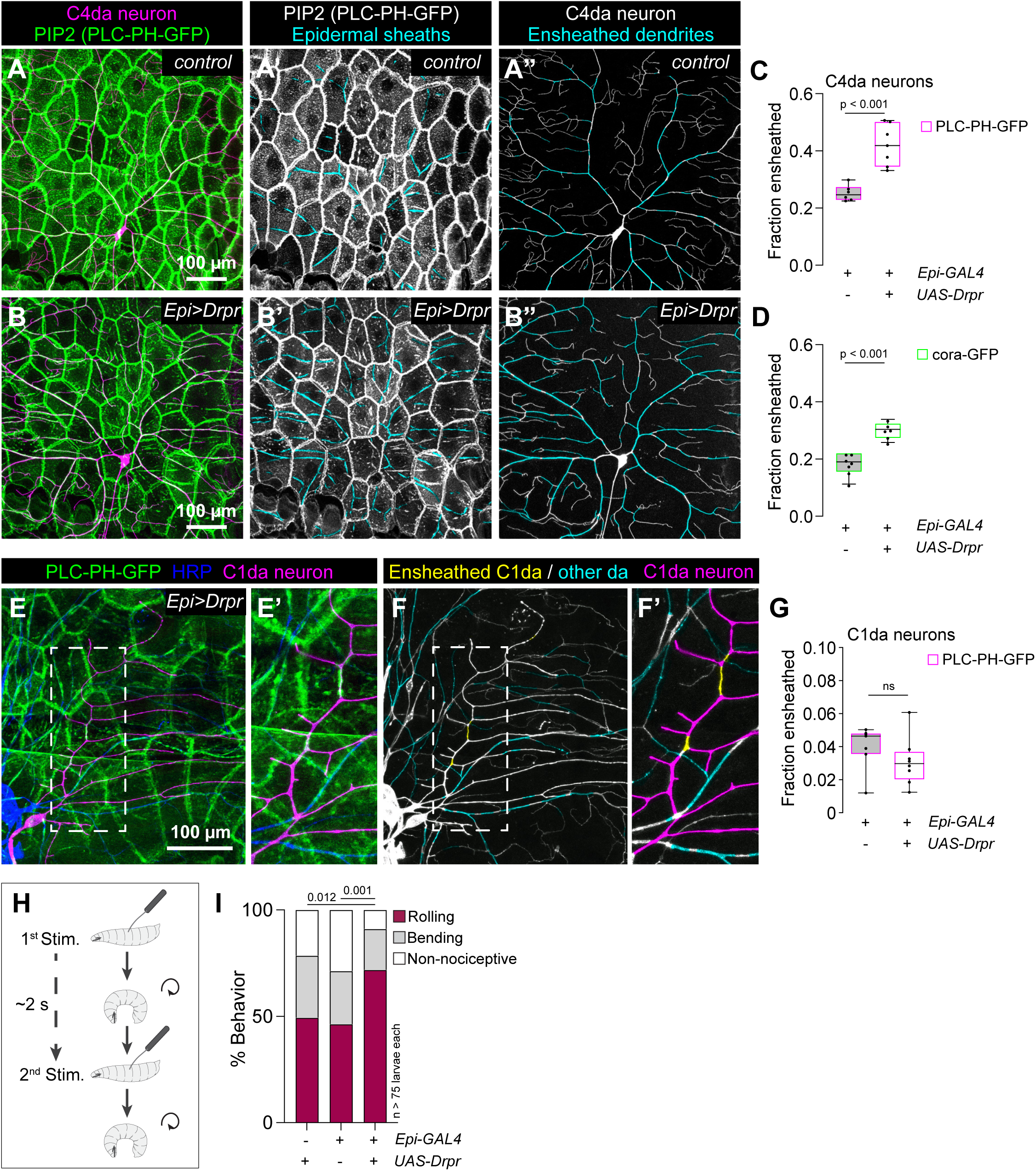
Drpr regulates the specificity of dendrite ensheathment. (A-C) Overexpressing *Drpr* in epidermal cells induces excess ensheathment. (A-B) Maximum intensity projections of confocal stacks show the distribution of C4da dendrites (magenta) and the PIP2 reporter PLC^δ^-PH-GFP to label epidermal sheaths (green) in an effector-only control larva (A) or a larva overexpressing *UAS-Drpr-I* in epidermal cells (B). Montages depict the distribution of epidermal sheaths (pseudocolored cyan) in a dorsal hemisegment (A’ and B’) and the extent of ensheathment (A” and B”) in representative neurons from control larvae and larvae overexpressing *UAS-Drpr-I* in epidermal cells. (C-D) Quantification of ensheathment defects. Plot depicts the proportion of C4da dendrite arbors that are ensheathed in larvae of the indicated genotypes. Sheaths were identified by labeling of epidermal membrane stretches adjacent to C4da dendrites (*ppk-CD4-tdTomato*) by (C) the PIP2 reporter PLC-PH-GFP or (D) or cora-GFP. n = 7 neurons per genotype, and *P<0.05 compared to wild-type controls, Mann Whitney test. (E-G) Overexpressing *Drpr* in epidermal cells does not induce ensheathment of C1da dendrites, which normally exhibit limited ensheathment. (E) Composite image depicts antibody labeling of sensory neurons (anti-HRP) and the epidermal sheath reporter PLC^δ^-PH-GFP (anti-GFP) with the C1da neuron ddaE pseudocolored magenta. (F) Montage depicts the distribution of PIP2-positive epidermal sheaths in cyan and C1da dendrites in magenta. Dendrites of C1da neurons exhibit minimal ensheathment (yellow). (G) Plot depicts the fraction of C1da dendritic arbors ensheathed by PLC-PH-GFP-positive epidermal membrane domains. N = 8 neurons each genotype. No significant difference was detected between the groups using a Mann-Whitney test. (H-I) Overexpressing *Drpr* in epidermal cells triggers nociceptive sensitization to mechanical stimuli. (H) Schematic depicting the behavioral paradigm. Larvae were presented with two consecutive 50 mN mechanical stimuli spaced by a 5 sec recovery and the response to the second stimulus is shown in (I). Epidermal *Drpr* overexpression yielded a significant increase in nocifensive behavioral responses to mechanical stimulus. n > 75 larvae for each genotype. P values, Chi-square test. <u>Experimental Genotypes:</u> (A-G) *<u>control</u>: w^1118^; ppk-CD4-tdTomato^4A^ / +; A58-GAL4, UAS-PLC^δ^-PH-GFP, ppk-CD4-tdTomato^10A^ / +* *<u>Epi>Drpr</u>*: *w^1118^; ppk-CD4-tdTomato^4A^ / UAS-Drpr-I; A58-GAL4, UAS-PLC^δ^-PH-GFP, ppk-CD4-tdTomato^10A^ / +* (H-I) *<u>Effector control</u>: w^1118^; UAS-Drpr-I / +* *<u>Driver control</u>: w^1118^;; R38F11-GAL4 / +* *<u>Experimental</u>: w^1118^; UAS-Drpr-I / +; R38F11-GAL4 / +*

Harsh touch activates C4da neurons to elicit stereotyped nocifensive rolling responses (Zhong et al., 2010), and treatments that reduce epidermal dendrite ensheathment attenuate nocifensive responses to mechanical stimuli (Jiang *et al*., 2019). To determine whether this effect on mechanical nociceptive sensitivity is bidirectional, we investigated effects of epidermal *Drpr* overexpression on larval mechanonociception. For these studies we used a paradigm that generates robust/reproducible nocifensive output and assayed for the frequency of larval rolling and bending behaviors (Tenedini *et al*., 2025). Briefly, we presented larvae with two successive mechanical stimuli and assayed for nocifensive behavioral responses to the second stimulus (Figure 5G). In driver and effector controls we observed comparable responses, with 49% and 47% of larvae exhibiting nocifensive rolling responses to the second of two consecutive 50 mN stimuli (Figure 5H). In contrast, larvae overexpressing *Drpr* in epidermal cells exhibited a significantly higher frequency of rolling responses (71%) and nocifensive responses overall.

### PS exposure promotes dendrite ensheathment

Phagocytosis of degenerating neurites and apopotic cells by Drpr involves recognition of extracellular PS (MacDonald *et al*., 2006; Doherty *et al*., 2009; Lu *et al*., 2017); indeed, PS-exposure on neurites is sufficient to trigger epidermal engulfment (Sapar *et al*., 2018; Ji *et al*., 2023). We therefore hypothesized that molecular recognition of dendrites for epidermal ensheathment by Drpr is mediated by PS exposure on dendrites. To test this model, we first assayed for PS accumulation at sites of ensheathment using a genetically encoded sensor for extracellular PS, a secreted form of the PS-binding protein AnnexinV tagged with the fluorescent protein mScarlet (AnnV-mScarlet) (Ji *et al*., 2023) (Figure 6A). Consistent with our model, we found that PLC-PH-GFP-positive epidermal membrane stretches were labeled by secreted AnnV-mScarlet (Figure 6, A and B).

**Figure 6.**
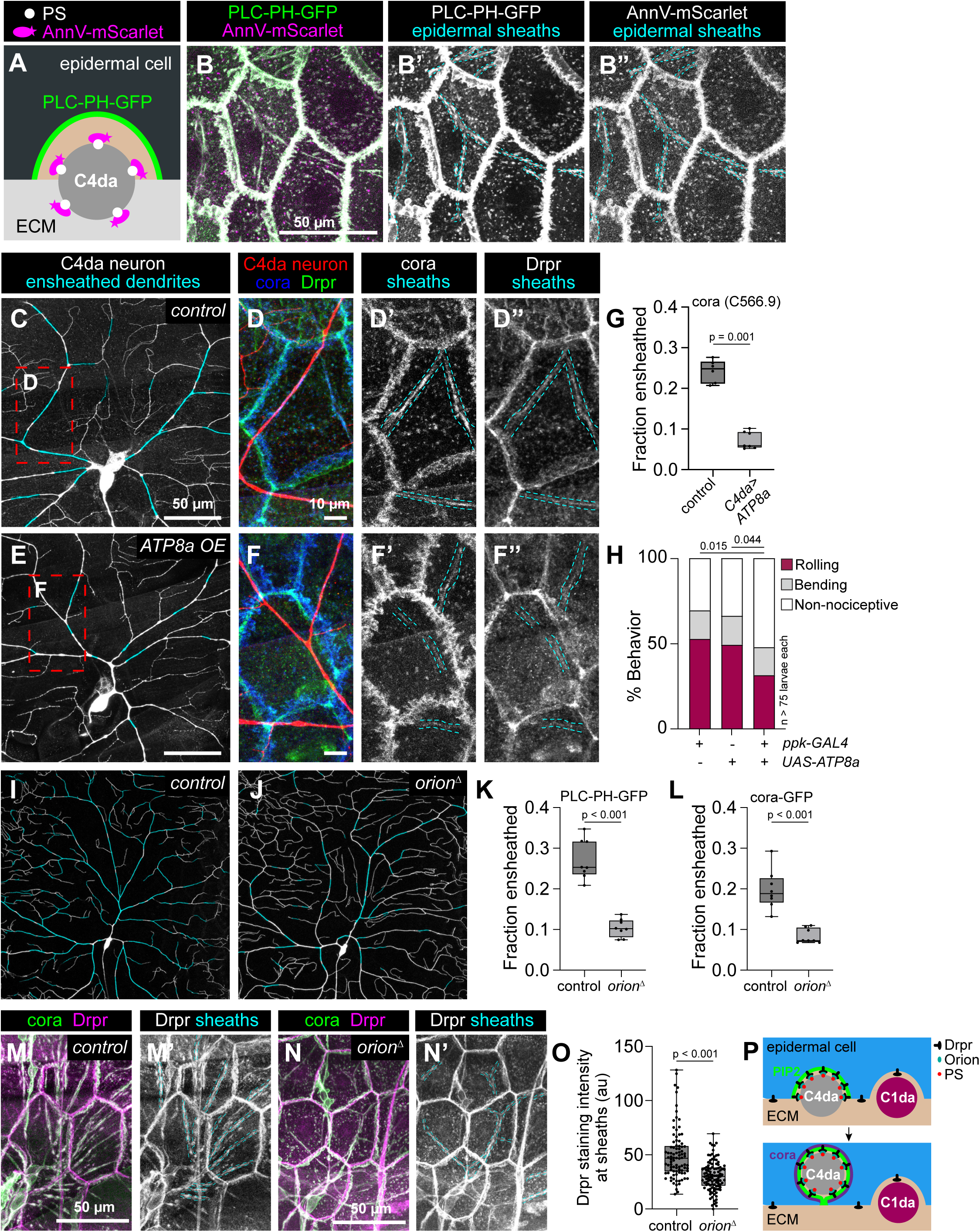
Phosphatidyl serine (PS) exposure likely promotes Drpr-dependent epidermal dendrite ensheathment. (A-B) Epidermal sheaths are co-labeled by a marker of externalized PS. (A) Schematic depicting reporters used to label epidermal sheaths (PLC-PH-GFP) and PS exposed on C4da dendrites. (B) Maximum intensity projection depicts dual labeling of the larval epidermis with markers for epidermal PIP2 in green and externalized PS in magenta. Both markers extensively accumulate at junctional domains and sites of dendrite ensheathment. Single channel images depict distribution of PLC-PH-GFP (B’) and AnnV-mScarlet labeling (B”) with epidermal sheaths marked with cyan hatched lines. (C-G) Overexpressing *ATP8a* in C4da neurons to limit PS exposure prevents epidermal ensheathment. Montages depict distribution of cora-positive sheaths in a single C4da neuron from a representative control (C) or a larva overexpressing the phospholipid flippase *ATP8a* selectively in C4da neurons (E). Red hatched boxes mark the regions shown in maximum intensity projections (D and F), which depict immunofluorescence labeling with antibodies to GFP (C4da neurons), cora (D’ and F’) and Drpr (D” and F”). Cyan hatched lines mark the position of epidermal sheaths in single channel images. (G) Neuronal *ATP8a* overexpression blocks ensheathment. Plot depicts the proportion of C4da arbors contained within cora+ sheaths in larvae of the indicated genotypes. n = 8 neurons for each genotype. P < 0.05, Mann-Whitney test. (H) Neuronal *ATP8a* overexpression reduces mechanical nociceptive sensitivity. Plot depicts behavioral responses of larvae of the indicated genotypes to the second of two successive 50 mN mechanical stimuli. n > 75 larvae per genotype, P values, Chi-square test. (I-L) The PS-bridging molecule Orion is required for ensheathment. Montages depict the distribution of PIP-2-positive epidermal sheaths on C4da dendritic arbors of representative neurons from (I) control (*orion* heterozygote) or (J) *orion* mutant larvae. Homozygous *orion* mutant larvae exhbit significant reductions in epidermal ensheathment compared to heterozgyous controls. Plots depict the proportion of C4da arbors ensheathed by (K) PLC-PH-GFP-positive and (L) cora(1-383)-GFP-positive epidermal membrane domains. n = 8 neurons for each genotype/reporter combination. P values, Mann-Whitney test. (M-O) *Orion* promotes Drpr recruitment to epidermal sheaths. Maximum intensity projections depict distribution of cora (to label epidermal sheaths) and Drpr immunoreactivity in representative control (M) and *orion* mutant (N) larvae. Single channel images (M’ and N’) depict Drpr immunoreactivity with the position of epidermal sheaths marked by cyan hatched lines. (O) Plot depicts the intensity of Drpr immunoreactivity (background subtracted) at epidermal sheaths of control and orion mutant larvae. Each point represents the mean intensity of Drpr signal across an individual sheath, and measurements were taken from 8 neurons of each genotype. P value, Mann-Whitney test. (P) Model depicting contributions of Drpr-Orion-PS interactions in ensheathment. <u>Experimental Genotypes:</u> (A-B) *w^1118^;; A58-GAL4, UAS-PLC^δ^-PH-GFP / UAS-AnnV-mScarlet* (C-G) *<u>control</u>*: *w^1118^; ppk-CD4-tdGFP^1b^ / +* *<u>ATP8a OE</u>*: *w^1118^; ppk-GAL4, ppk-CD4-tdGFP^1b^ / UAS-ATP8a* (H) *<u>Effector control</u>: w^1118^; UAS-ATP8a / +* *<u>Driver control</u>*: *w^1118^;; ppk-GAL4 / +* *<u>Experimental</u>: w^1118^; UAS-ATP8a / +; ppk-GAL4 / +* (I-L) *<u>control</u>: orion^/1^/+; ppk-CD4-tdTomato^4A^ / +; A58-GAL4, UAS-PLC^δ^-PH-GFP, ppk-CD4-tdTomato^10A^ / +* *orion^/1^: orion^/1^; ppk-CD4-tdTomato^4A^ / +; A58-GAL4, UAS-PLC^δ^-PH-GFP, ppk-CD4-tdTomato^10A^ / +* (M-O) *<u>control</u>*: *orion^/1^/+* *orion^/1^: orion^/1^*

Next, we evaluated effects of blocking PS externalization on dendrite ensheathment and associated phenotypes. We reasoned that if PS exposure induces epidermal dendrite ensheathment, then preventing PS exposure should block sheath formation. The PS flippase ATP8A drives unidirectional PS translocation from the outer to the inner leaflet of the plasma membrane (Sapar *et al*., 2018), and we found that overexpressing *UAS-ATP8A* selectively in C4da neurons using *ppk-GAL4* yielded a significant reduction in epidermal sheaths (Figure 6, C to G). Furthermore, similar to other treatments that reduce epidermal ensheathment (Jiang *et al*., 2019), we found that overexpressing *ATP8A* in C4da neurons led to a significant reduction in nocifensive responses to mechanical stimuli (Figure 6H). Finally, recognition of damaged dendrites for phagocytosis by epidermal cells requires the secreted chemokine-like PS-bridging molecule Orion (Boulanger *et al*., 2021; Ji *et al*., 2023), and we found that *orion* mutation significantly reduced the overall extent of epidermal ensheathment (Figure 6, I to L, and Supplemental Figure 4) as well as Drpr recruitment to epidermal sheaths (Figure 6, M to O). Altogether, these results support the model that epidermal Drpr, together with the secreted PS-bridging molecule Orion, recognizes PS externalized on dendrites to promote ensheathment (Figure 6P).

## Discussion

Multiple intercellular recognition events govern SSN morphogenesis in the skin. Here, we identified mediators of the selective ensheathment of certain SSN neurites by epidermal cells. Specifically, we found that the nimrod receptor Drpr functions in epidermal cells to drive ensheathment of SSN dendrites labeled by extracellular PS. Our studies define several features of this evolutionarily conserved intercellular interaction. First, multiple receptors likely promote sheath formation, as *Drpr* inactivation reduces but does not eliminate ensheathment. Second, PS exposure on neurites appears to be a primary signal for ensheathment: reporters for extracellular PS are enriched at sites of ensheathment, and overexpressing PS flippases in C4da neurons or inactivation of the PS bridging molecule Orion blocks ensheathment. Third, the level of neuronal signals is not the only determinant of the extent of ensheathment: overexpressing *Drpr* in epidermal cells increased the extent of ensheathment without altering substrate specificity, suggesting that neuronal signals are not limiting. Fourth, epidermal phagocytic receptors appear to regulate multiple distinct aspects of ensheathment. We found that RNAi of the nimrod-family receptor *Nimb4* led to an increase in sheaths that were not innervated by dendrites, suggestive of exuberant sheath growth or failures in sheath disassembly following neurite retraction. *Nimb4* encodes a secreted protein that binds apoptotic cells and promotes phagosome maturation following engulfment (Petrignani *et al*., 2021), hence it will be intriguing to determine whether sheath disassembly involves autophagy. Finally, our studies suggest that the extent of epidermal ensheathment dynamically tunes nociceptor sensitivity to mechanical stimuli; whether it likewise modulates responses to other noxious inputs and/or other sensory modalities remains to be determined.

What are the molecular mediators of dendrite recognition that drive ensheathment? Prior studies established that L1CAM/Neuroglian (Nrg) regulates dendrite positioning of SSN dendrites to potentiate ensheathment, with epidermal cells and SSNs expressing different Nrg isoforms that are capable of interacting (Yamamoto *et al*., 2006; Yang *et al*., 2019). However, Nrg tethers both ensheathed and unensheathed dendrites to the epidermis (Yang *et al*., 2019), hence additional recognition events are required to account for the selectivity of ensheathment. Our studies demonstrate that recognition of exposed PS on neurites by epidermal Drpr is one key determinant of ensheathment, but additional factors likely contribute as well. *Drpr* inactivation caused a ∼50% reduction in ensheathment, and we found no sign of persistent Drpr protein in *Drpr* mutants during a time window when sheaths continued to form, hence other epidermal receptors likely mediate recognition events that promote ensheathment. Neuronal overexpression of the flippase ATP8A to block PS exposure led to an almost complete loss of ensheathment, suggesting that externalized PS may be a requisite signal for recognition events that drive ensheathment. Hence, additional epidermal receptors could directly or indirectly recognize externalized PS on dendrites to promote ensheathment, and several lines of evidence point towards Integrins as candidate ensheathment receptors. Selective knockdown of Integrin expression in epidermal cells blocks dendrite ensheathment and phenocopies ensheathment-defective dendrite morphogenesis phenotypes (Jiang *et al*., 2014). Integrins are heterodimeric receptors that contribute to PS-mediated phagocytosis in both professional and non-professional phagocytes (Finnemann and Silverstein, 2001; Sexton et al., 2001; Hsieh et al., 2012; Flannagan et al., 2014), though Integrins do not directly bind PS. Instead, Integrins are linked indirectly to PS, for example by the opsonin MFG-E8 and PS receptor TIM4 (Akakura *et al*., 2004; Nandrot *et al*., 2007). *Drosophila* Integrins contribute to phagocytic engulfment in multiple contexts, including Drpr-independent functions in hemocytes and Drpr-dependent apoptotic engulfment of dying germline cells by follicular epithelium (Nagaosa *et al*., 2011; Etchegaray *et al*., 2012; Meehan *et al*., 2015). The latter involves Integrin complexes containing βPS/Myospheroid (Mys), the predominan β-Integin subunit found in the epidermis, and αPS3/Scab, which is required for the morphogenetic movements during would healing in the larval epidermis (Park *et al*., 2018). Hence, future studies are warranted to explore contributions of epidermal Integrins to epidermal sheath formation.

Although PS exposure is commonly associated with apoptosis, PS exposure does not always trigger phagocytic recognition (Sakuragi and Nagata, 2023), and indeed our studies indicate that PS functions in a context-dependent manner to trigger epidermal phagocytic engulfment or ensheathment of sensory neurites. What accounts for these different outcomes? Studies of non-apoptotic functions of PS provide several possible mechanisms that could influence epidermal responses to exposed PS. Healthy mouse lymphoma cells that constitutively expose PS on the cell surface by virtue of an activating mutation in the TMEM16F scramblase are not bound by circulating macrophages, but treatment with pro-apoptotic signals triggers macrophage engulfment (Segawa *et al*., 2011). These results suggest that alteration of the PS that is exposed and/or signals in addition to PS confer information about context in these cells. PS exposed during apoptosis is often oxidized (Kagan *et al*., 2003) and some PS receptors exhibit heightened affinity to oxidized PS (Borisenko *et al*., 2004). Likewise, some receptors recognize different levels of exposed PS to trigger different outcomes, typified by the PS receptor T cell immunoglobulin and mucin-domain-containing molecule 4 (Tim4) which triggers apoptotic removal of high-PS exposing cells and removal of activated T-cells that expose lower levels of PS (Miyanishi *et al*., 2007). Mechanistically, Tim4 has a single PS-binding pocket as well as a series of basic residues that contribute weaker ionic interactions with membrane PS lipids, providing sensitivity to surface PS density (Tietjen *et al*., 2014). PS recognition by Drpr during phagocyting removal of injured dendrites relies on the extracellular PS bridging molecule Orion, whose levels can modulate epidermal sensitivity to PS (Ji *et al*., 2023). Orion is likewise required for epidermal dendrite ensheathment, which occurs on the same developmental timescale as phagocytosis of injured dendrites, hence factors other than Orion are likely required to differentially tune PS sensitivity of engulfing epidermal cells to discriminate between ensheathment and phagocytic engulfment. Finally, in some cases cells express signals that prevent apoptotic engulfment and/or PS recognition. For example, the cell surface molecule CD47 binds its receptor SIRPA on macrophages to protect from engulfment, with CD47 activating SIRPA at the phagocytic synapse where it inhibits Integrin activation (Gardai *et al*., 2005; Morrissey *et al*., 2020). Although *Drosophila* lacks identifiable CD47 homologs, an analogous mechanism could modulate the signaling output to PS engagement. Intriguingly, microglia which function as the primary phagocytes in the central nervous system partially envelop neurons during prion infection (Makarava *et al*., 2024). Hence, it will be intriguing to determine whether similar mechanisms govern the decisions of epidermal cells and microglia to phagocytose or envelop neurons.

What is the functional significance of epidermal dendrite ensheathment? Prior studies demonstrated that ensheathment stabilizes dendrite branches in *Drosophila* C4da neurons, and our work corroborates these findings. Whether epidermal sheaths similarly stabilize terminal arbors of unmyelinated C fibers remains to be determined, but in this context the reduction in intraepidermal nerve fiber (IENF) density that is commonly reduced in neuropathic conditions is noteworthy (Pittenger *et al*., 2005; Liu *et al*., 2014; Ghasemi and Rajabally, 2020). Our studies additionally show that the extent of epidermal ensheathment tunes sensitivity of nociceptors to mechanical stimuli. Mechanistically, several phenomena may contribute to this relationship between anatomical and functional coupling. Mature epidermal sheaths appear to form autotypic junctions, hence ensheathed neurites may be exposed to a distinctive extracellular environment that impacts excitability. Likewise, the ensheathed neurites may exhibit enhanced ionic coupling to epidermal cells (Jefferys, 1995; Anastassiou and Koch, 2015). Connexin43 plaques are present at epidermal sheaths in mammals (Erbacher *et al*., 2024), hence direct gap junctional coupling or hemi-channel release may provide a mechanism for preferential coupling of epidermal cells to ensheathed neurons. This finding is particularly noteworthy given prior studies demonstrating that keratinocyte stimulation triggers ATP release that may occur through gap junctions (Barr *et al*., 2013; Takada *et al*., 2014; Moehring *et al*., 2018). Finally, epidermal ensheathment may influence the transduction of mechanical inputs, similar the effect of dendrite intercalation at intercellular junctions which enhances mechanical sensitivity of C4da neurons in a Piezo-dependent fashion (Luedke *et al*., 2024).

## Materials and Methods

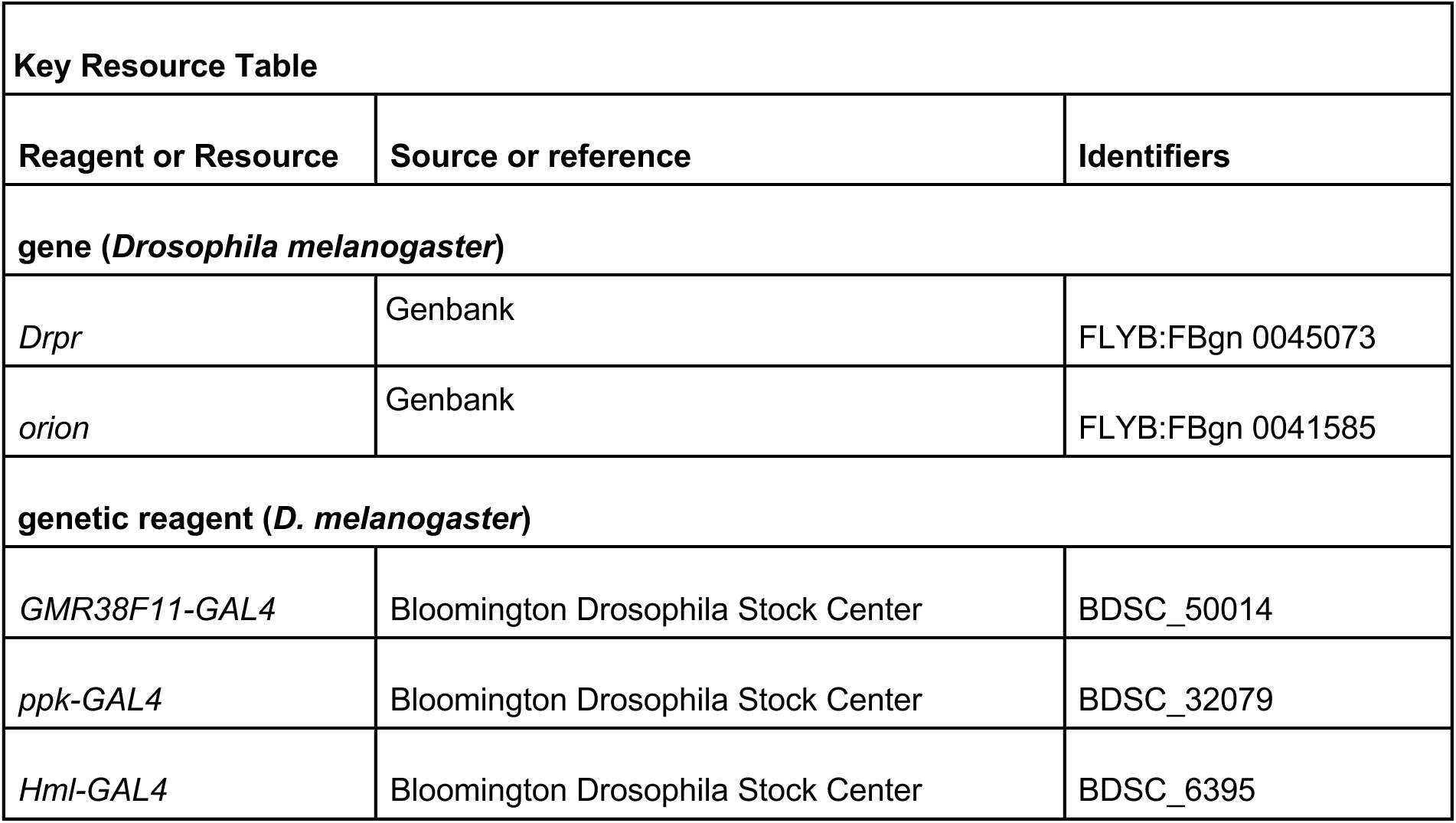
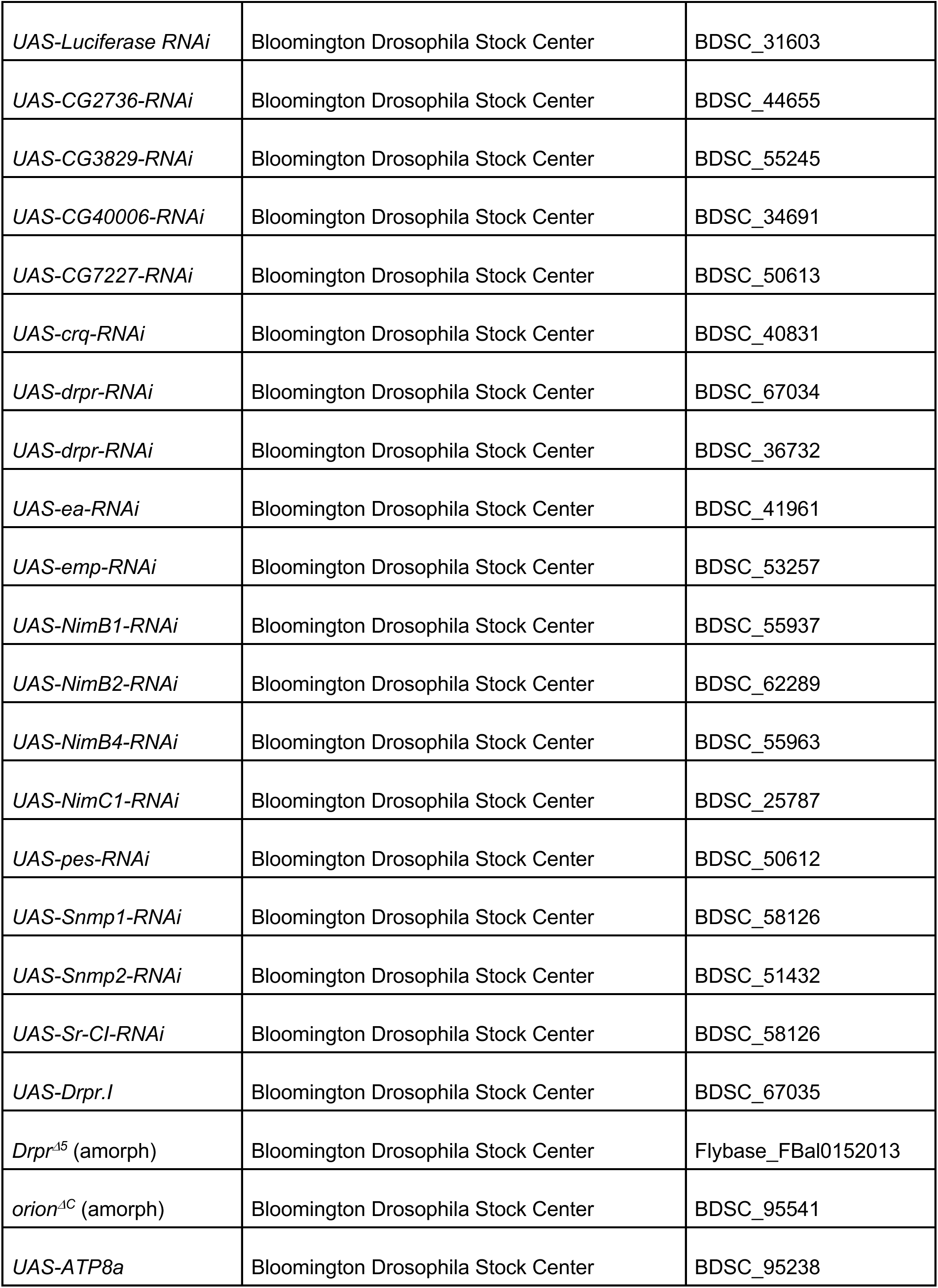
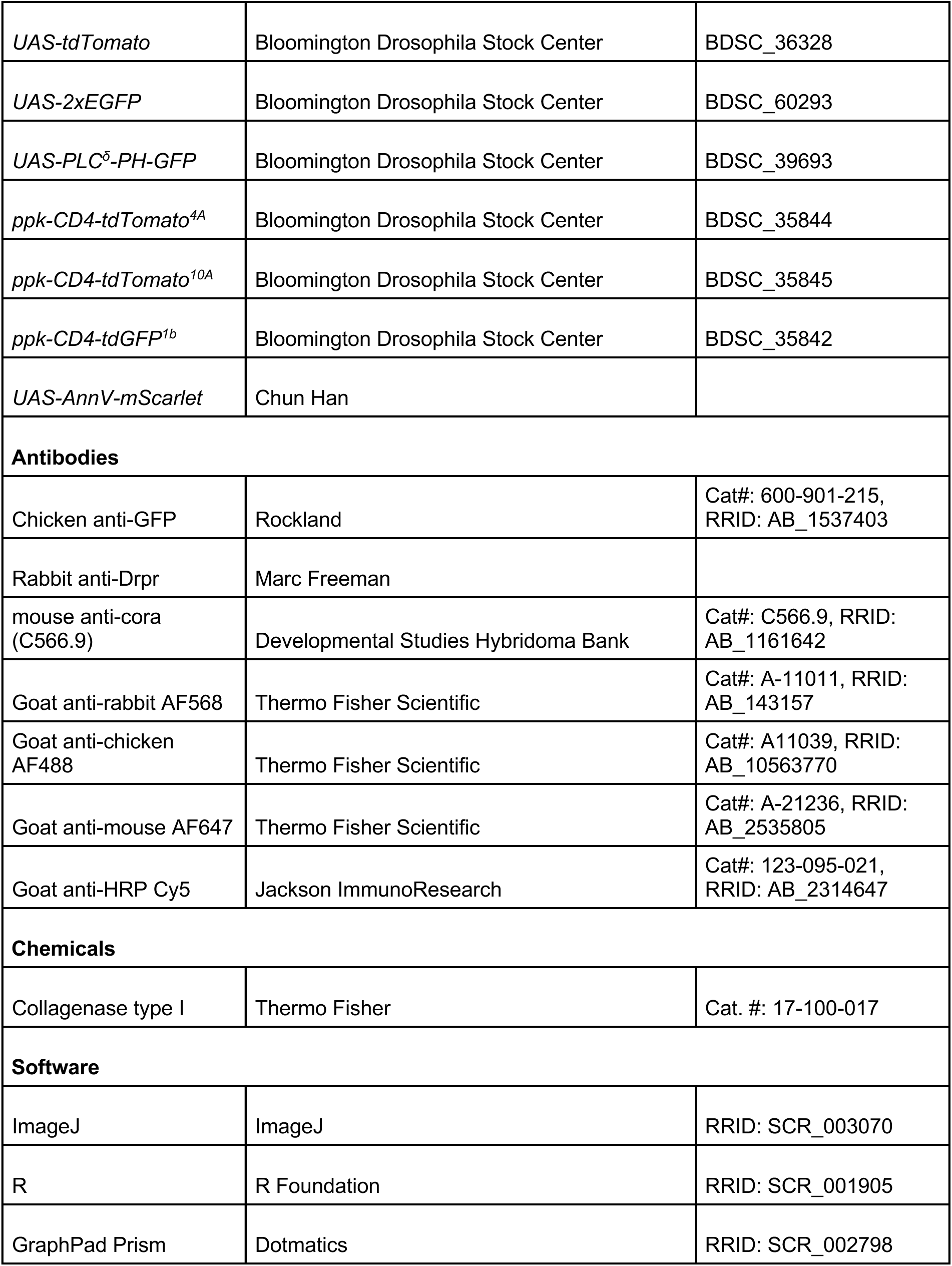

### Genetics

#### Drosophila strains

Flies were maintained on standard cornmeal-molasses-agar media and reared at 25° C under 12 h alternating light-dark cycles. A complete list of alleles used in this study is provided in the Key Resources Table; experimental genotypes are listed in figure legends.

### Microscopy

#### Live confocal imaging

Live single larvae were mounted in 90% glycerol under a coverslip and imaged on a Leica SP5 confocal microscope using a 20x 0.8 NA or 40x 1.25 NA lens. Larvae subject to time-lapse microscopy were recovered between imaging sessions to plates containing standard cornmeal-molasses-agar media.

#### Immunostaining

Third instar larvae were pinned on a sylgard plate, filleted along the ventral midline, and pinned open. After removing intestines, fat bodies, imaginal discs and ventral nerve cord, fillets were fixed in PBS with 4% PFA for 15 min at room temperature, washed 4 times for 5 min each in PBS with 0.3% Tx-100 (PBS-Tx), blocked for 1 h in PBS-Tx + 5% normal goat serum, and incubated in primary antibody overnight at 4° C. Samples were washed 4 times for 5 min each in PBS-Tx, incubated in secondary antibody for 4 h at room temperature, washed 4 times for 5 min each in PBS-Tx, and stored in PBS prior to imaging. Antibody dilutions were as follows: chicken anti-GFP (Aves Labs #GFP-1020, 1:500), mouse anti-coracle (DSHB, C566.9 supernatant, 1:25), Rabbit anti-Drpr (Logan *et al*., 2012) (1:500), HRP-Cy5 (Jackson Immunoresearch, 1:100), Goat anti-Chicken Alexa488 (Thermofisher A-11039, 1:200), Goat anti-rabbit Alexa 488 (Thermofisher A-11034, 1:200), Goat anti-Mouse Alexa555 (Thermofisher A-28180, 1:200), Goat anti-rabbit Alexa555 (Thermofisher A-21428, 1:200), Goat anti-mouse AF647 (Thermofisher A-21236, 1:200).

### Image analysis

#### Morphometric analysis of epidermal sheaths

Epidermal ensheathment of dendrites was measured from manual traces generated from composite images (neuronal marker + epidermal sheath marker) in Fiji (Schindelin *et al*., 2012). Ensheathed stretches (Jiang *et al*., 2019) were identified via co-localization with the epidermal PIP2 marker PLC^δ^-PH-GFP or cora immunoreactivity and manually traced in Fiji. Sheath width values represent means from measurements taken at 1 µm intervals along the entire sheath length. C4da neurons were identified by co-labeling with C4da-specific reporters, and C1da neurons were identified by their stereotyped morphology using anti-HRP labeling. Quantitative analysis of the extent of ensheathment (Figures 1, 2, 5, 6) was conducted on da neurons in dorsal hemi-segments A3 and A4 using a field of view of 387.5 x 387.5 mm, which captures the majority of the dendritic arbor of a single C4da neuron.

#### Drpr accumulation at epidermal sheaths

Image stacks were collected under conditions to limit signal saturation, using identical settings for matched control/experimental specimens. To evaluate Drpr accumulation at epidermal sheaths, ROIs were drawn around individual sheaths labeled by PLC-PH-GFP or anti-cora immunoreactivity and mean fluorescence intensity of anti-Drpr immunoreactivity was measured within the ROI. Background intensity of anti-Drpr immunoreactivity was measured in an identically sized ROI adjacent to the sheath for background subtraction and in each epidermal cell across the entire specimen (excluding labeling at sheaths and junctional domains). Sheaths were scored Drpr+ if background-subtracted signal intensity exceeded 2x the standard deviation of epidermal background values.

#### Morphometric analysis of C1da and C4da dendrites

Features of dendrite morphology, including dendrite branch number, dendrite length, and dendrite crossing number were measured in Fiji (Schindelin *et al*., 2012) using the Simple Neurite Tracer plugin (Longair *et al*., 2011).

### RNA-Seq analysis of epidermal cells

#### RNA isolation for RNA-Seq

RNA-Seq analysis of hemocytes was performed concurrently with previously described RNA-Seq analysis of larval epidermal cells and C4da neurons (Yoshino *et al*., 2025). Larvae with cytoplasmic GFP expressed in hemocytes (*UAS-2x-EGFP/+; hml-GAL4/+*) were microdissected and dissociated in collagenase type I (Fisher 17-100-017) into single cell suspensions as previously described (Williams *et al*., 2016), with the addition of 1% BSA to the dissociation mix. After dissociation, cells were transferred to a new 35 mm petri dish with 1 mL 50% Schneider’s media / 50% PBS supplemented with 1% BSA. Under a fluorescent stereoscope, individual fluorescent cells were manually aspirated with a glass pipette into PBS with 0.5% BSA, and then serially transferred until isolated from all cellular debris. Twenty hemocytes per sample were aspirated together, transferred to a mini-well containing 3ul lysis solution (0.2 % Triton X-100 in water with 2 U / µL RNAse Inhibitor), lysed by pipetting up and down several times, transferred to a microtube, and stored at -80° C. For the picked cells, 2.3 µL of lysis solution was used as input for library preparation.

#### RNA-Seq library preparation

RNA-Seq libraries were prepared from the picked cells following the Smart-Seq2 protocol for full length transcriptomes (Picelli *et al*., 2014). To minimize batch effects, primers, enzymes, and buffers were all used from the same lots for all libraries. Libraries were multiplexed, pooled, and purified using AMPure XP beads, quality was checked on an Agilent TapeStation, and libraries were sequenced as 51-bp single end reads on a HiSeq4000 at the UCSF Center for Advanced Technology.

#### RNA-Seq data analysis

Reads were demultiplexed with CASAVA (Illumina) and read quality was assessed using FastQC (https://www.bioinformatics.babraham.ac.uk/) and MultiQC (Ewels *et al*., 2016). Reads containing adapters were removed using Cutadapt version 2.4 (Martin, 2011)(Martin, 2011) and reads were mapped to the *D. melanogaster* transcriptome, FlyBase genome release 6.29, using Kallisto version 0.46.0 (Bray *et al*., 2016) with default parameters. AA samples were removed from further analysis for poor quality, including low read depth (< 500,000 reads), and low mapping rates (< 80%). Raw sequencing reads and gene expression estimates are available in the NCBI Sequence Read Archive (SRA) and in the Gene Expression Omnibus (GEO) under the following accession numbers: GSE284380 for epidermal cells; SUB15311573 for hemocytes.

### Experimental Design and Statistical Analysis

For each experimental assay control populations were sampled to estimate appropriate sample numbers to allow detection of ∼33% differences in means with 80% power over a 95% confidence interval. Datasets were tested for normality using Shapiro-Wilks goodness-of-fit tests. Details of statistical analysis including treatment groups, sample numbers, statistical tests, controls for multiple comparisons, p-values and q-values are provided in Supplemental Methods.

## Acknowledgements

This work was supported by funding from the National Institutes of Health (NIH NINDS R01 NS076614 and NINDS R21NS125795 to J.Z.P), awards from the Weill Neurohub to J.Z.P and F.M.T, an award from the Scan Design Foundation to J.Z.P., and startup funds from UW to J.Z.P. Fly stocks were obtained from the Bloomington Drosophila Stock Center, which was supported by funding from the NIH (NIHP40OD018537). Antibodies obtained from the Developmental Studies Hybridoma bank, created by the NICHD of the NIH and maintained at The University of Iowa, were used in this study. We also thank Marc Freeman for sharing anti-Drpr antiserum, Chun Han for sharing fly stocks used in this study, Kazuo Emoto, Peter Soba, and the Parrish lab for helpful discussions, and Jeff Rasmussen for critical reading of the manuscript.

## Author Contributions

Acquisition of imaging data: A.P. and J.Z.P.

RNA-seq analysis: C.Y.

Acquisition of behavioral data: F.T.

Analysis and interpretation of data: imaging data, C.Y. and J.Z.P; RNA-seq data, C.Y. and J.Z.P

Drafting the article: J.Z.P.

## Conflict of Interest

The authors declare that they have no conflict of interest.

---

**Supplemental Figure 1.**
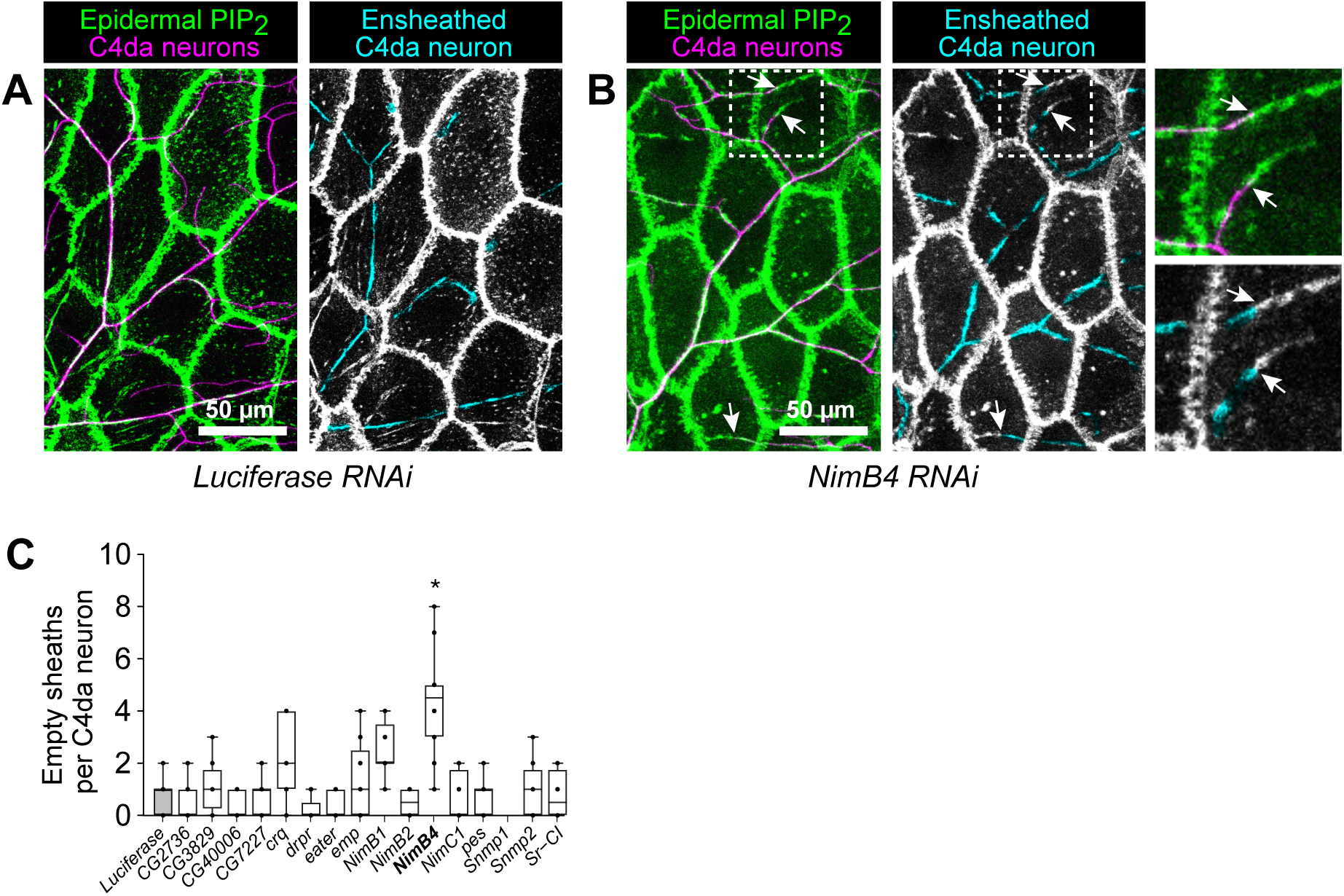
Related to. **Figure 1**. Multiple phagocytic receptors regulate different aspects of epidermal dendrite ensheathment. (A-B) The nimrod receptor gene *NimB4* is required for proper coupling of dendrites and epidermal sheaths. (A) Maximium intensity projections show dual color labeling of epidermal PIP2 and C4da dendrites and montages depict the distribution of epidermal sheaths in cyan. *Nimb4 RNAi* larvae contained stretches of epidermal sheaths that were not invaded by dendrites (white arrowheads). Zoomed images depict two epidermal sheaths that are not completely invaded by C4da dendrites. (B) Plot depicts the number of epidermal sheaths per C4da neuron with stretches that lacked dendrite invasion. *P<0.05 compared to *Luciferase RNAi* control; one-way ANOVA with post-hoc Dunnett’s multiple comparisons test. <u>Experimental Genotypes</u>: *w^1118^; ppk-CD4-tdTomato^4A^ / +; A58-GAL4, UAS-PLC^δ^-PH-GFP, ppk-CD4-tdTomato^10A^ / UAS-RNAi transgene*

**Supplemental Figure 2.**
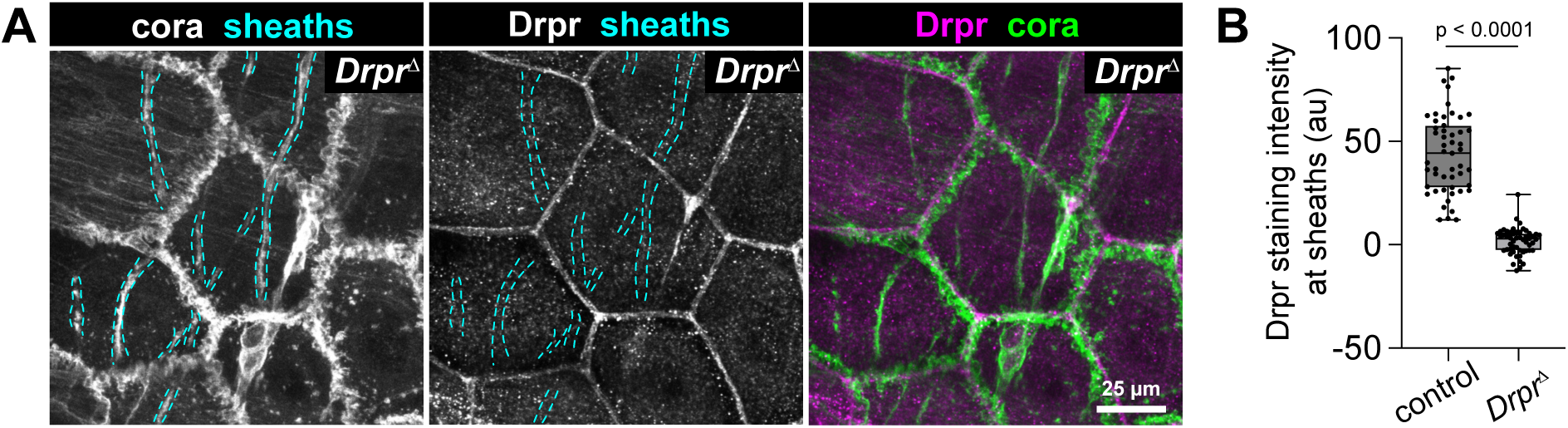
Specificity of anti-Drpr antibody labeling. Related to Figure 3. (A) Maximum intensity projections of confocal stacks depict anti-cora immunoreactivity, with epidermal sheaths marked by cyan hatched lines, anti-Drpr immunoreactivity, and merged image. Drpr immunoreactivity at sheaths is absent from *Drpr* mutant larvae as is the labeling of septate junctions that is seen in wild-type controls (see Figure 3). Signal remains in the apical junctional domain, suggesting that the polyclonal Drpr antibody additionally recognizes an antigen in a protein other than Drpr that localizes to this domain. (B) Signal intensity of anti-Drpr immunoreactivity at cora+ epidermal sheaths. Each point represents the mean signal intensity along the length of an entire sheath (with background staining along a parallel membrane domain 1 μm away from the sheath), and staining intensity at sheaths from 5 neurons was measured (n > 50 sheaths each). P value, Mann-Whitney test. <u>Experimental Genotypes</u>: (A) *w^1118^;; Drpr^/15^* (B) *w^1118^* (control), *w^1118^;; Drpr^/15^* (*Drpr*)

**Supplemental Figure 3.**
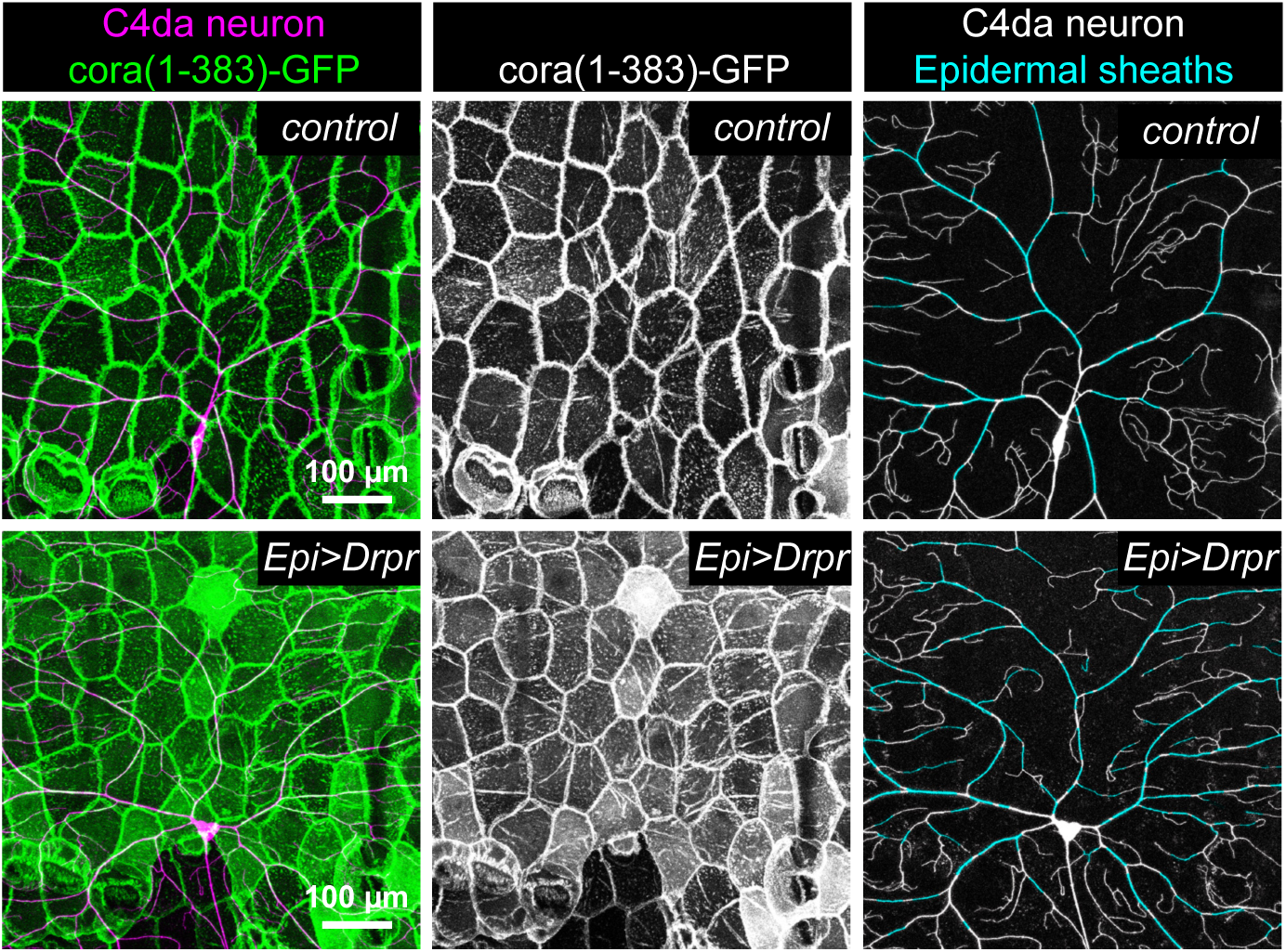
**Epidermal *Drpr* overexpression drives increased ensheathment**. Related to Figure 5. Dual labeling nociceptive C4da neurons (*ppk-CD4-tdTomato*) together with (A) the PIP2 marker PLC^δ^-PH-GFP or (B) the mature epidermal sheath marker cora(1-383)-GFP in representative control larvae or larvae overexpressing *Drpr* (*UAS-Drpr-I*) in epidermal cells. Maximum projections of confocal stacks show the distribution of C4da dendrites and the epidermal sheath marker (left) and the epidermal sheath marker alone (center) in a single dorsal hemisegment. The composite images (right) depict the distribution of epidermal sheaths along the C4da arbor. Genotypes: (A) *A58-GAL4, UAS-PLC^δ^-PH-GFP / +* (control) or *A58-GAL4, UAS-PLC^δ^-PH-GFP / UAS-Drpr-I*. (B) *A58-GAL4, UAS-cora(1-383)-GFP / +* (control) or *A58-GAL4, UAS-cora(1-383)-GFP / UAS-Drpr-I*.

**Supplemental Figure 4.**
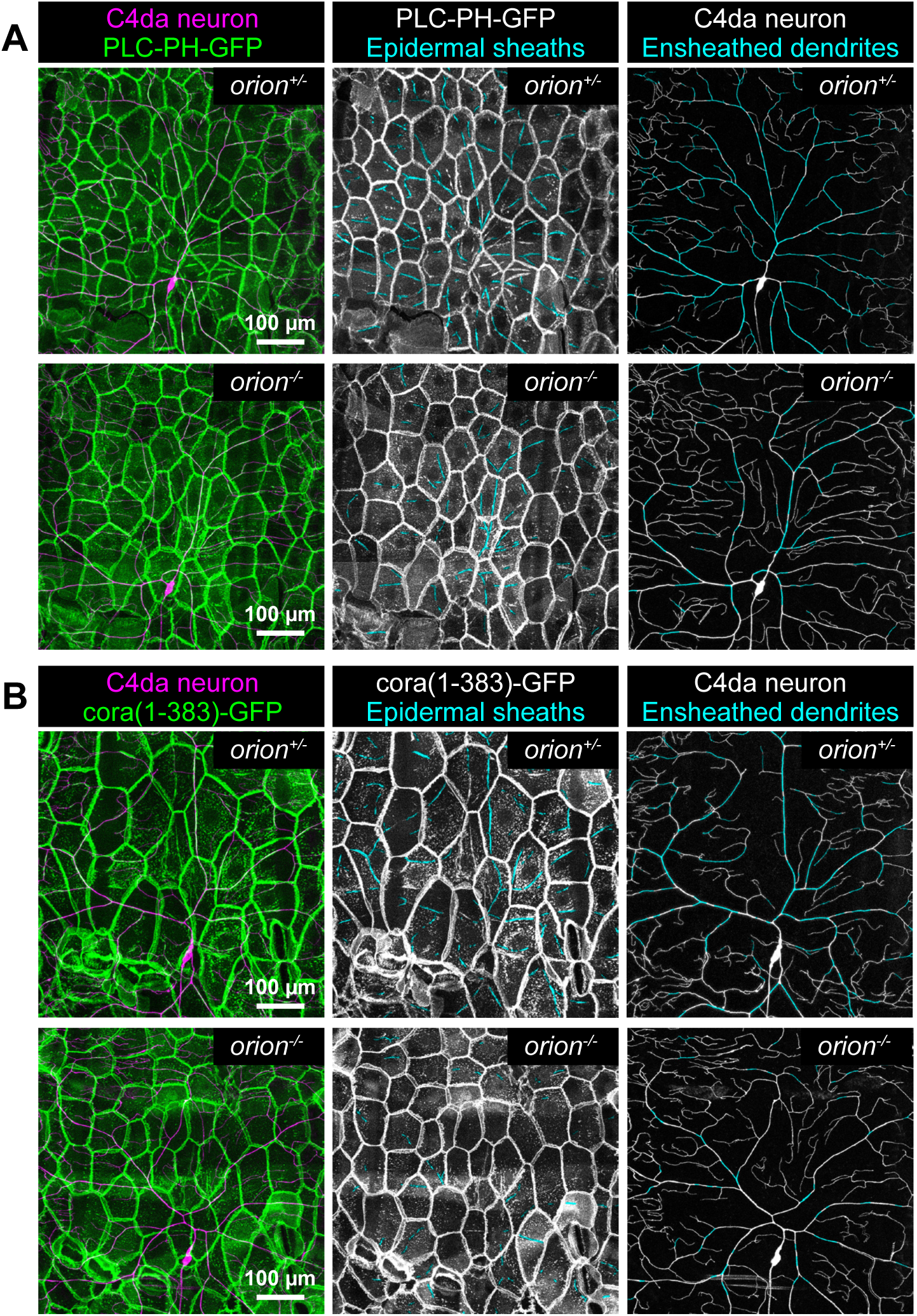
O***r***ion **mutation reduces formation of epidermal sheaths.** Related to Figure 6. Dual labeling nociceptive C4da neurons (*ppk-CD4-tdTomato*) together with (A) the PIP2 marker PLC^δ^-PH-GFP or (B) the mature epidermal sheath marker cora(1-383)-GFP in representative *orion* heterozygous (control) or homozygous mutant larvae. Maximum projections of confocal stacks show the distribution of C4da dendrites and the epidermal sheath marker (left) and the epidermal sheath marker alone (center) in a single dorsal hemisegment. The composite images (right) depict the distribution of epidermal sheaths along the C4da arbor. Genotypes: (A) *orion^/1^ / +; A58-GAL4, UAS-PLC^δ^-PH-GFP / +* (control) or *orion^/1^ / orion^/1^; A58-GAL4, UAS-PLC^δ^-PH-GFP / +*. (B) *orion^/1^ / +; A58-GAL4, UAS-cora(1-383)-GFP / +* (control) or *orion^/1^ / orion^/1^; A58-GAL4, UAS-cora(1-383)-GFP / +*.

**Fig. 1F:**
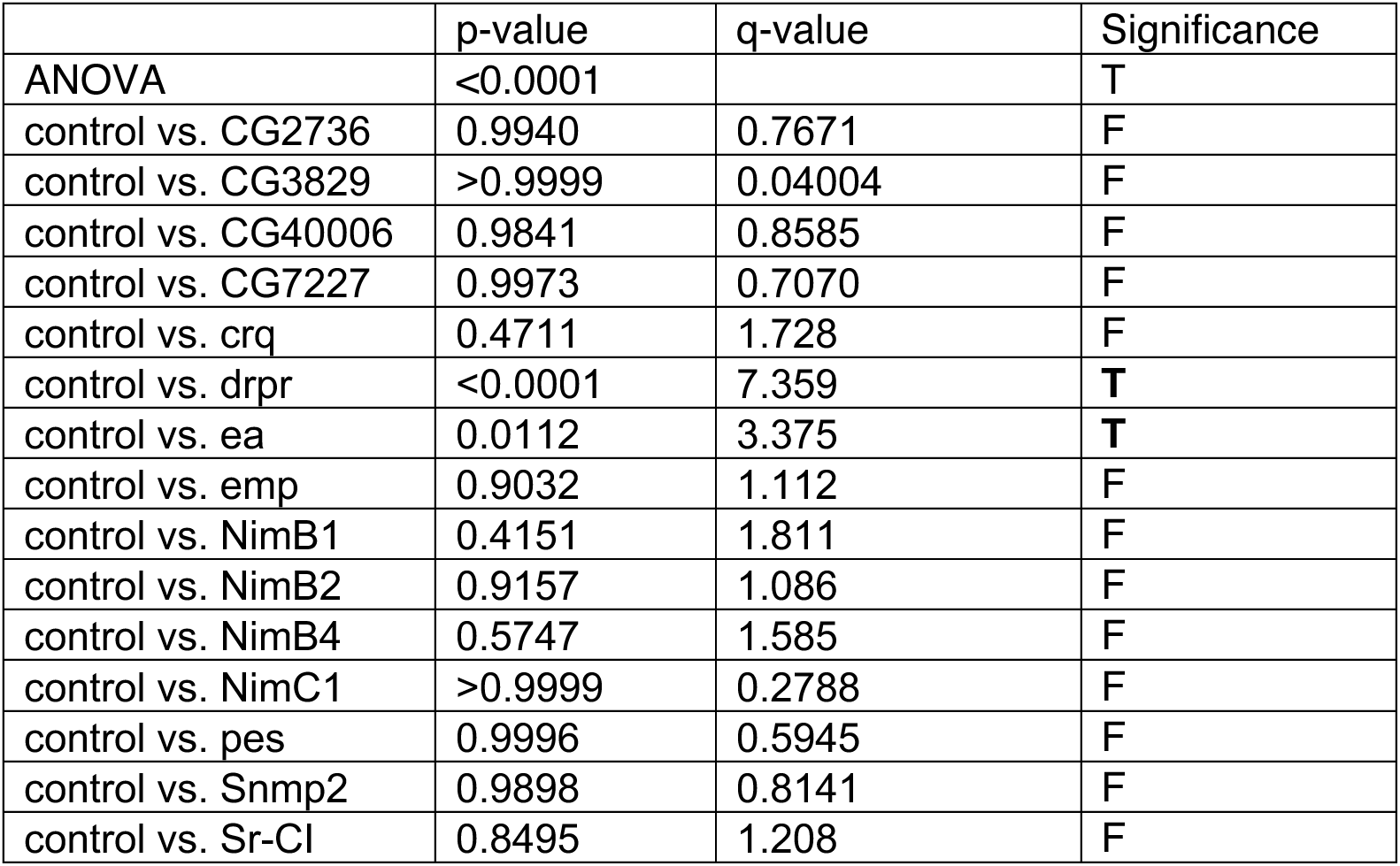
One-way ANOVA followed by Dunnett’s multiple comparisons test

**Figure 1G:**
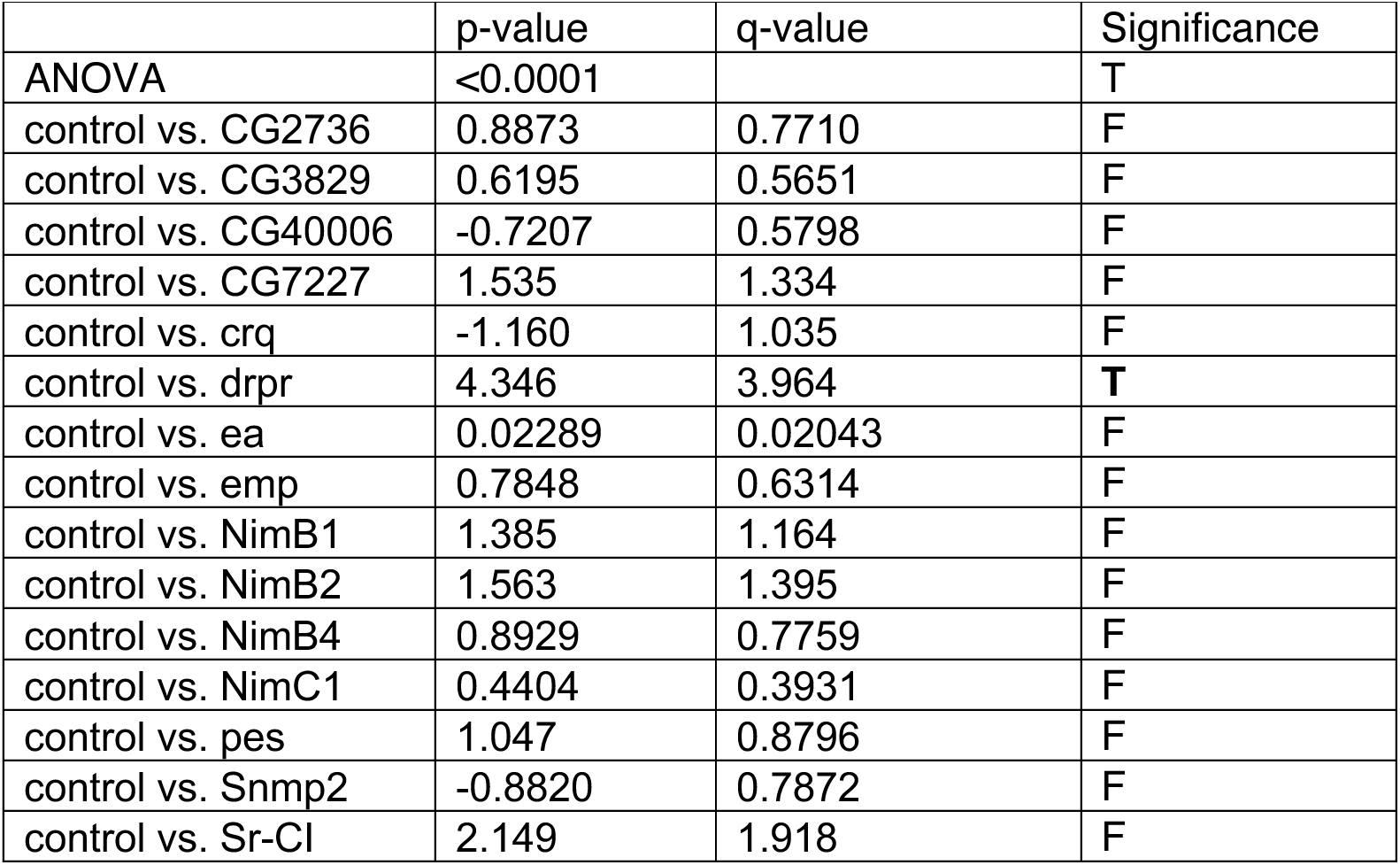
One-way ANOVA followed by Dunnett’s multiple comparisons test

**Figure 1H:**
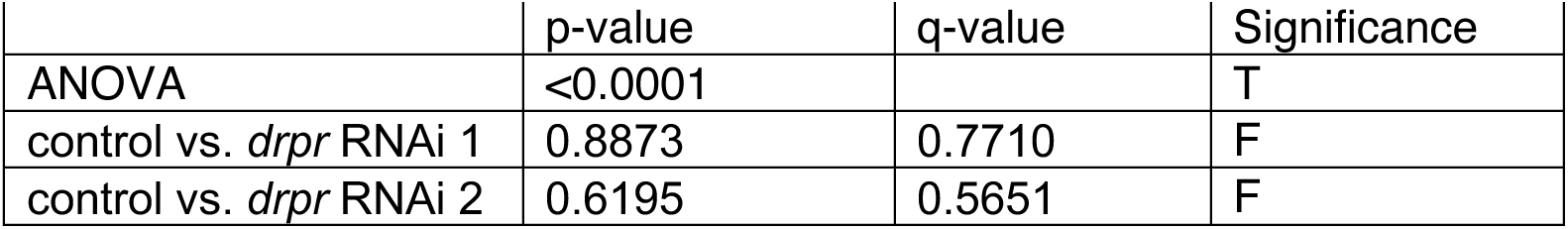
One-way ANOVA followed by Dunnett’s multiple comparisons test

**Figure 1I:**
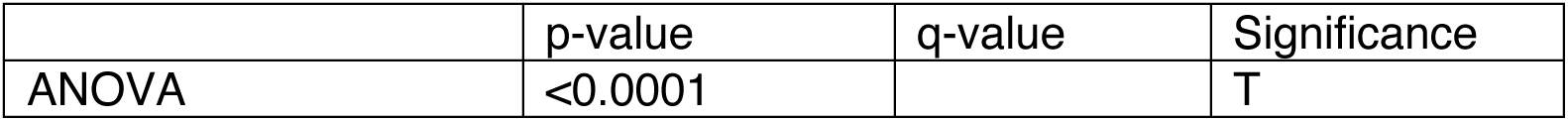
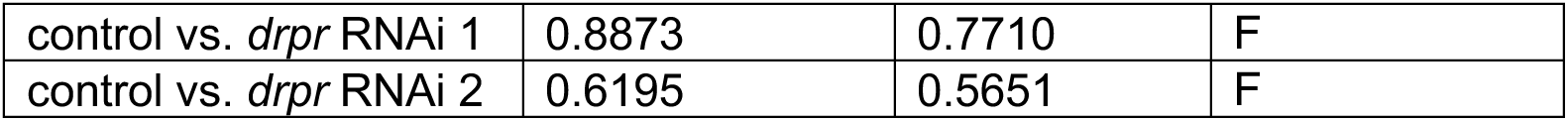
One-way ANOVA followed by Dunnett’s multiple comparisons test

**Figure 1J:**
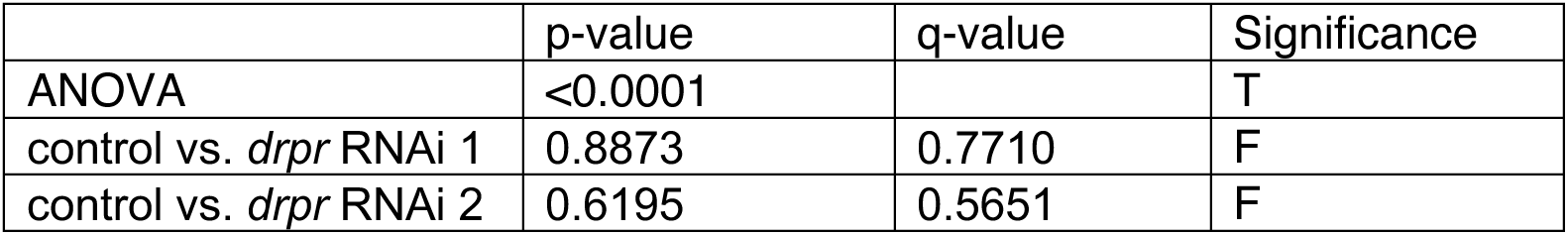
One-way ANOVA followed by Dunnett’s multiple comparisons test

**Figure 1K:**
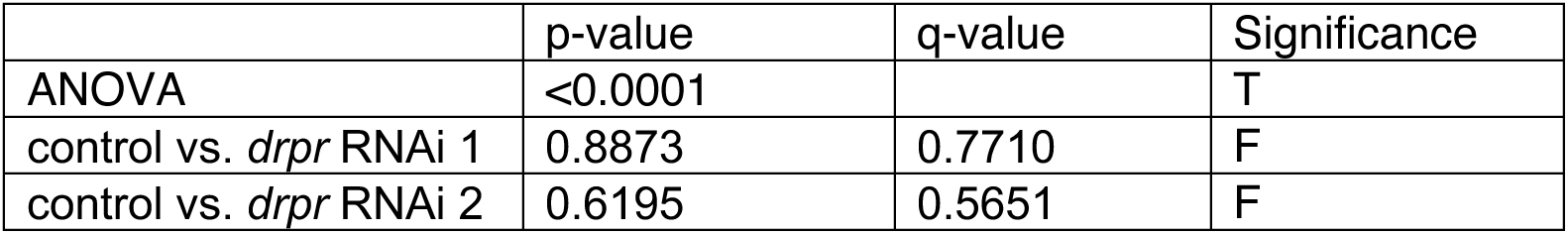
One-way ANOVA followed by Dunnett’s multiple comparisons test

**Figure 2C:**
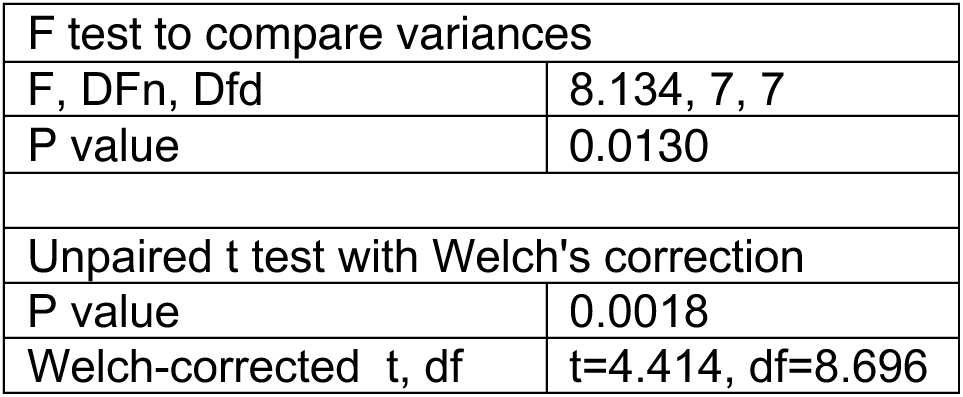
cora+ sheath number

**Figure 2D:**
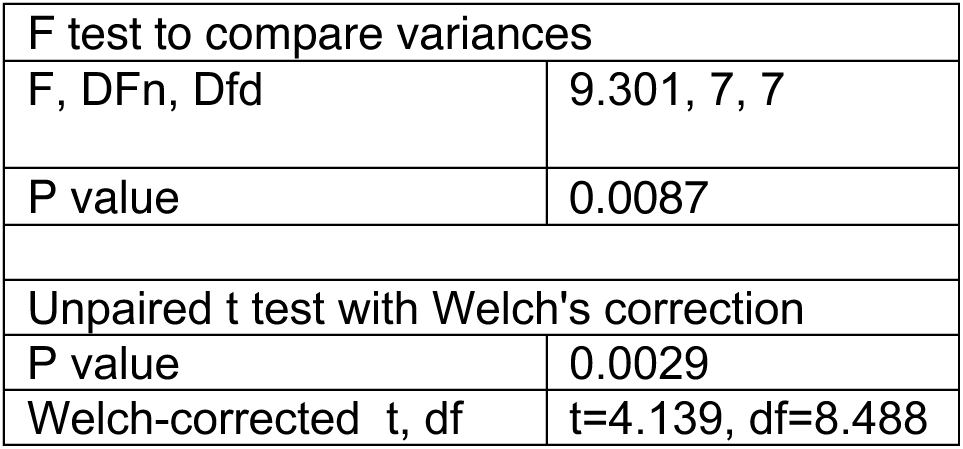
cora+ sheath density.

**Figure 2E:**
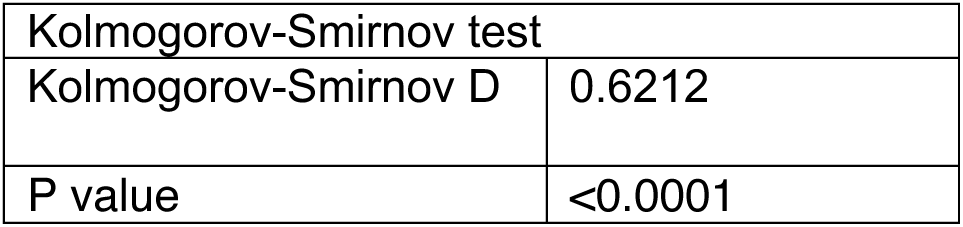
cora+ sheath width

**Fig. 2F:**
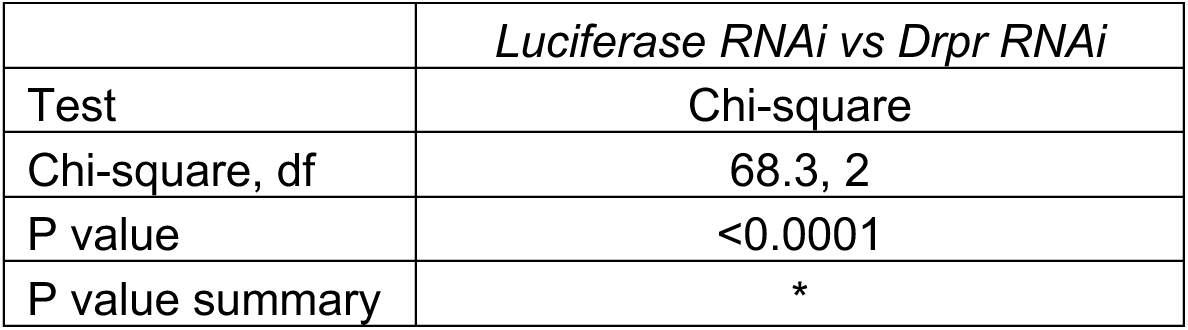
Sheath dynamics

**Figure 2G:**
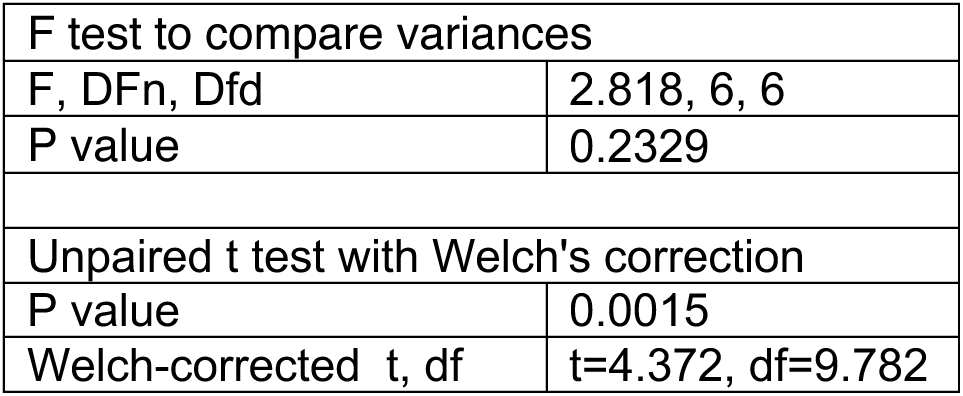
Change in sheath length

**Figure 4B:**
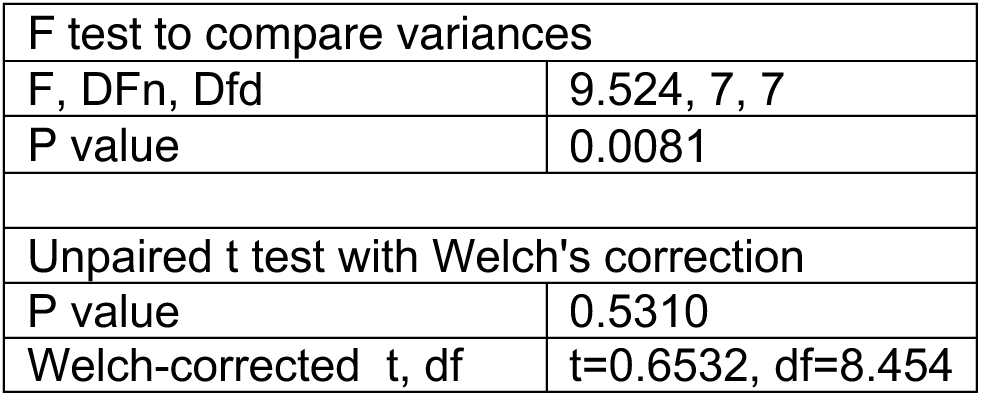
C1da dendrite length

**Figure 4C:**
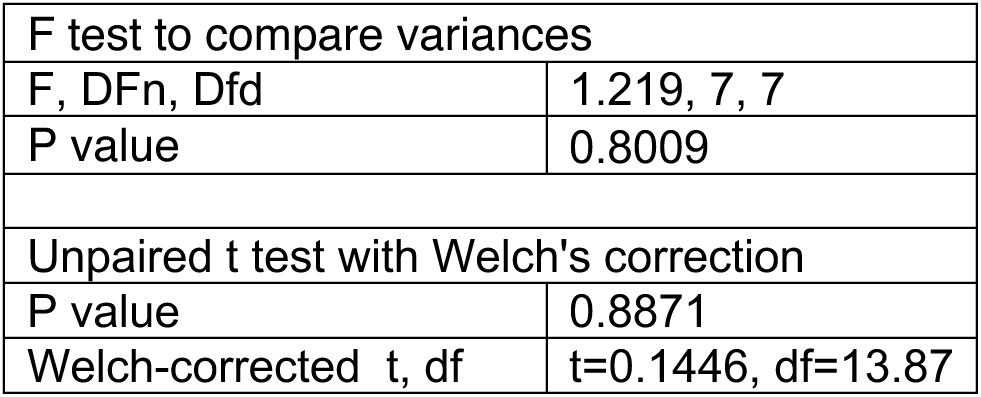
C1da dendrite branchpoints

**Figure 4E:**
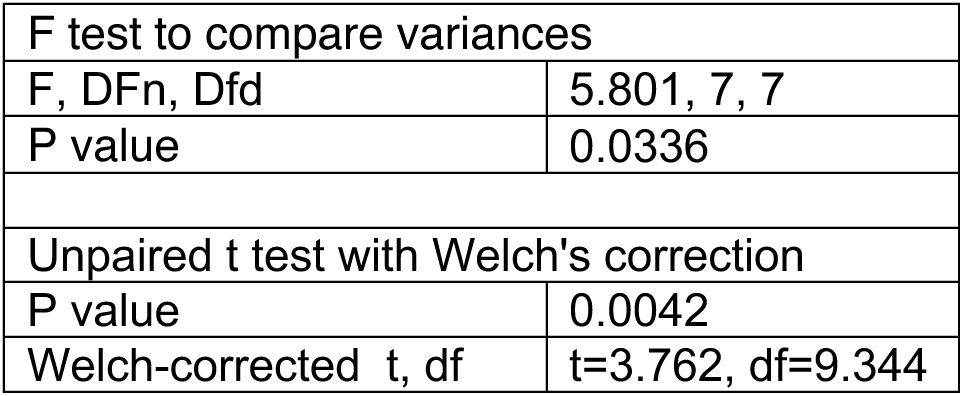
Normalized C4da dendrite length

**Figure 4F:**
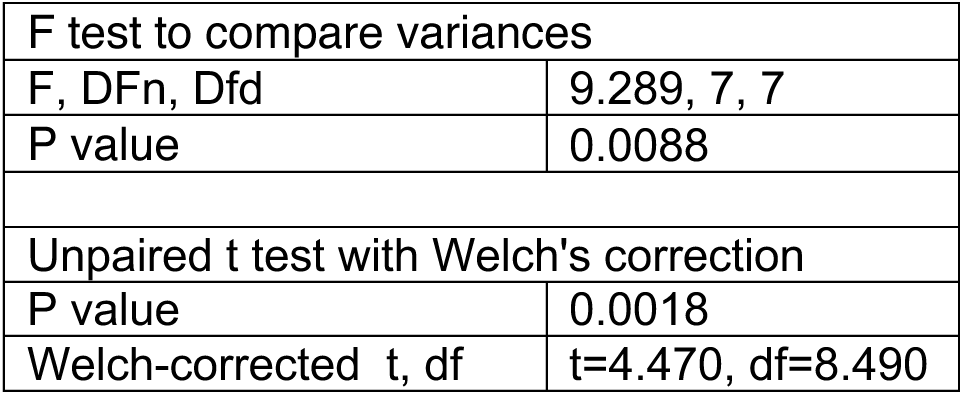
C4da dendrite terminal branch density

**Figure 4G:**
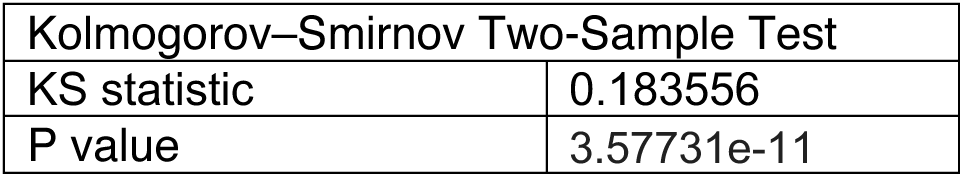
Distribution of dendrite terminal lengths

**Figure 5C:**
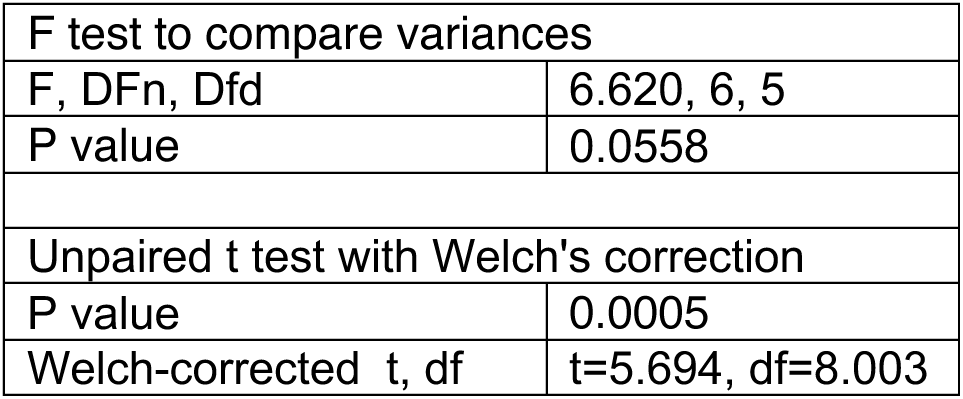
C4da dendrite ensheathment (PLC-PH-GFP)

**Figure 5D:**
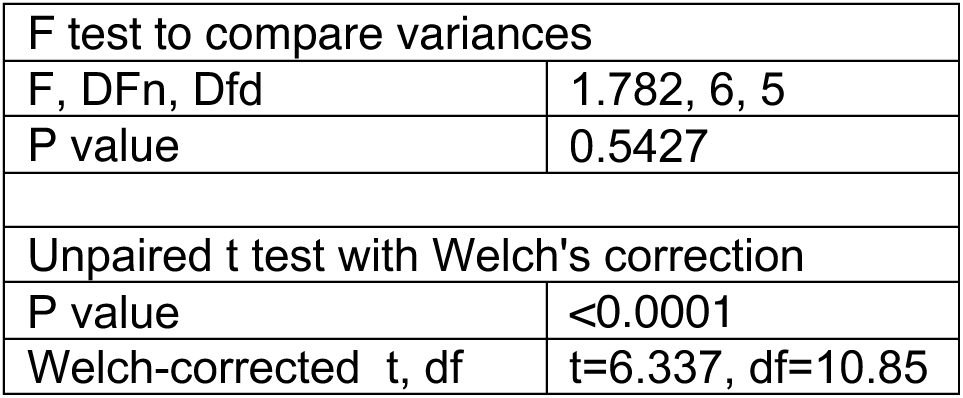
C4da dendrite ensheathment (cora-GFP)

**Figure 5G:**
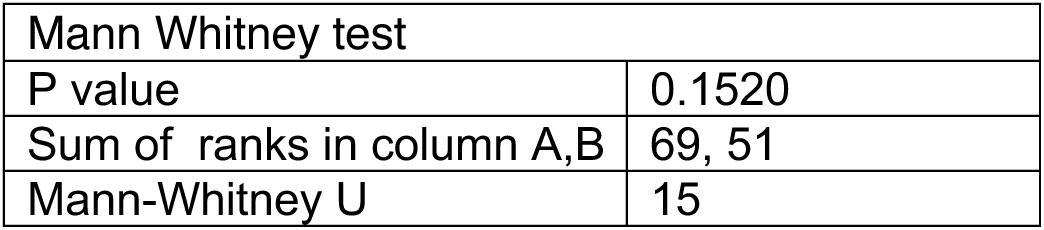
C1da dendrite ensheathment

**Figure 5I:**
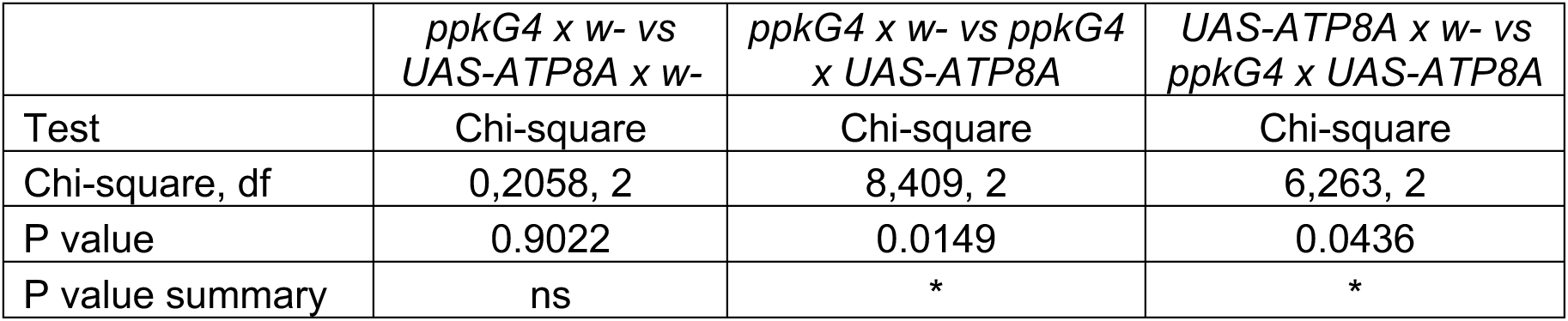
Nocifensive rolling responses.

**Figure 6G:**
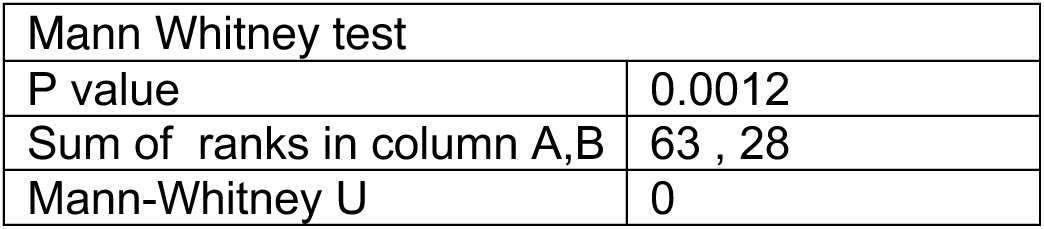
Fraction of C4da dendrites ensheathed

**Figure 6H:**
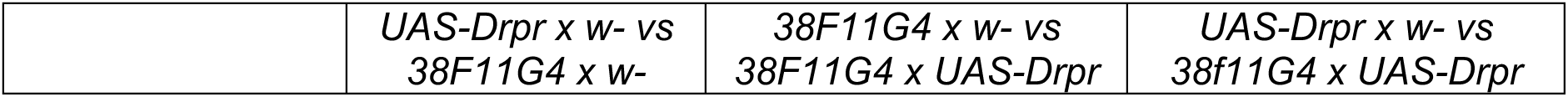
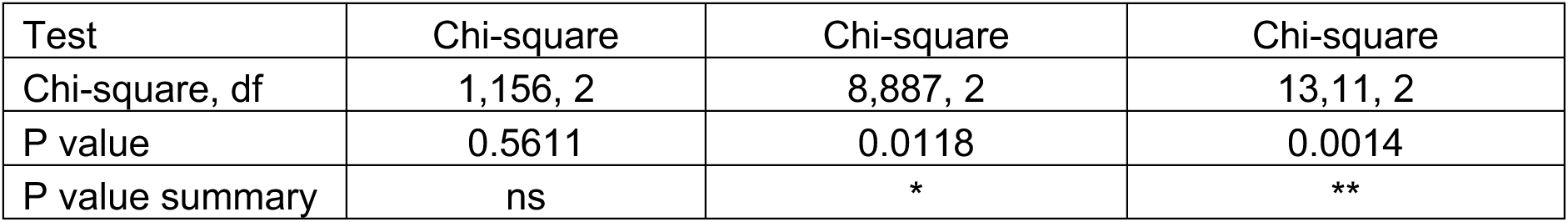
Nocifensive rolling responses.

**Figure 6K:**
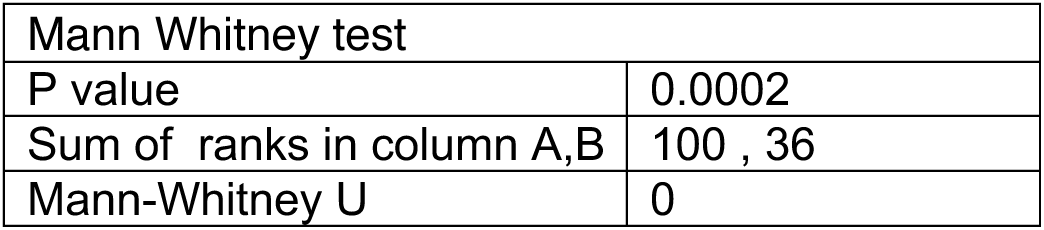
C4da dendrite ensheathment (PLC-PH-GFP)

**Figure 6L:**
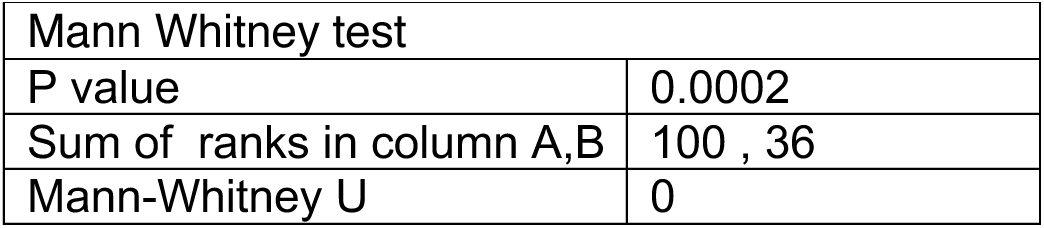
C4da dendrite ensheathment (cora-GFP)

**Figure 6O:**
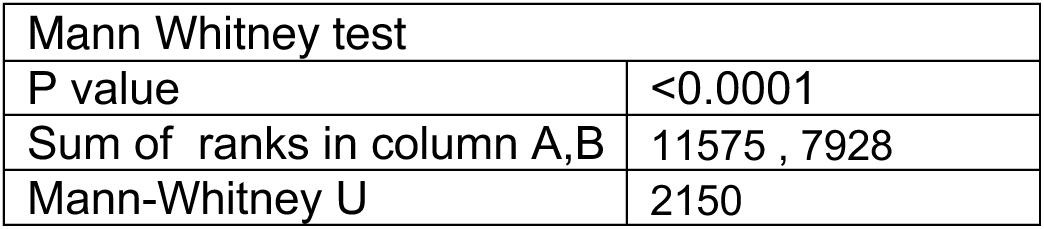
Drpr intensity at sheaths

**Supplemental Figure 1B.**
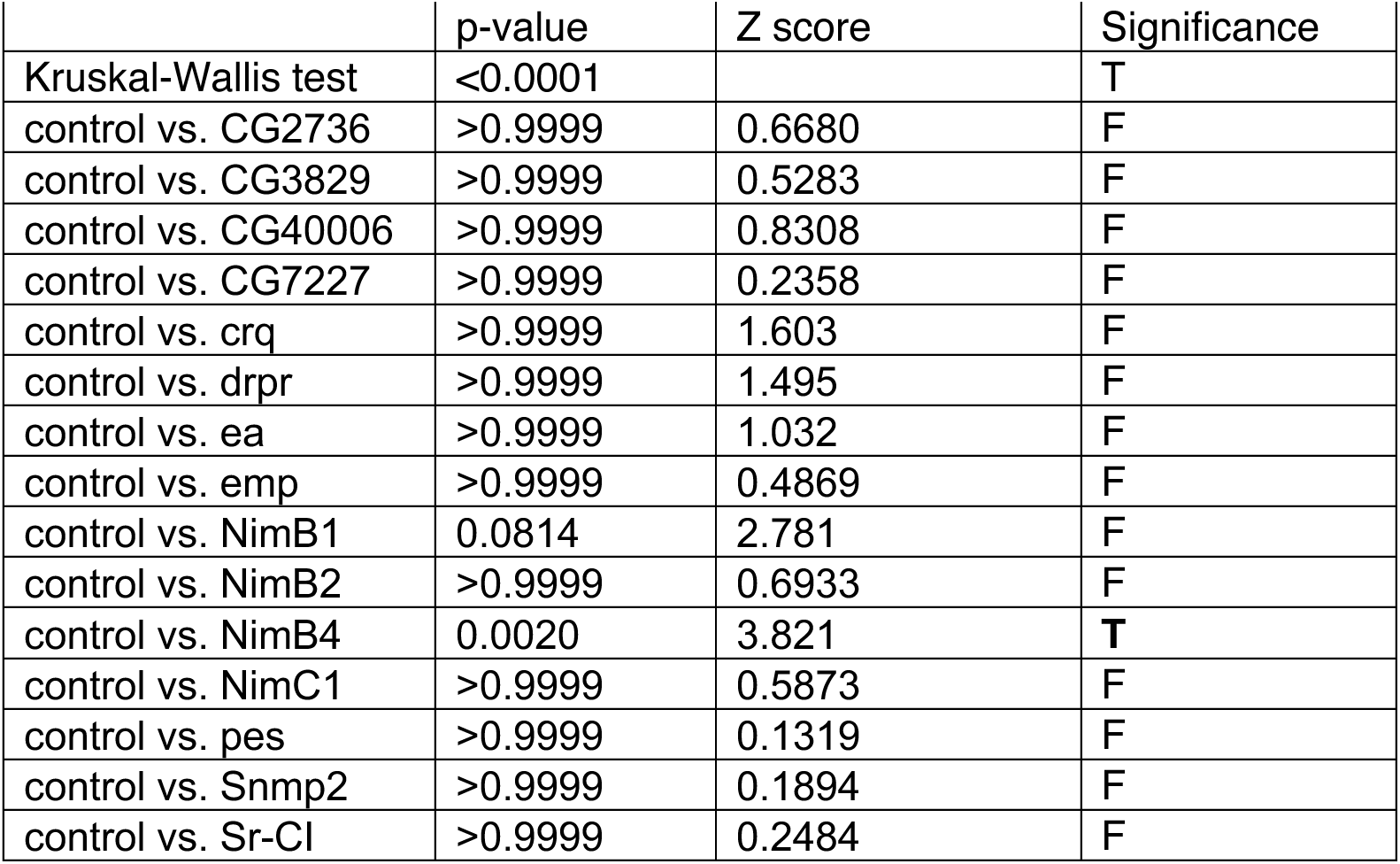
: Empty C4da sheaths. Kruskal-Wallis with post-hoc Dunn’s test

**Supplemental Figure 2B.**
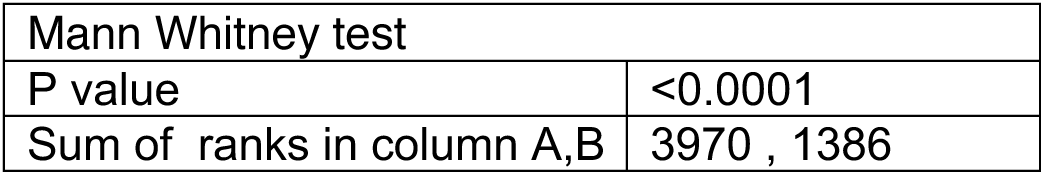
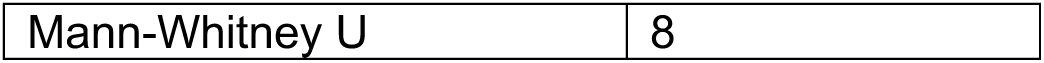
: Drpr intensity at sheaths in wt and *Drpr* mutant larvae.

## Notes

### Competing Interest Statement

The authors have declared no competing interest.

